# Reuniens transiently synchronizes memory networks at beta frequencies

**DOI:** 10.1101/2022.06.21.497087

**Authors:** Maanasa Jayachandran, Tatiana D. Viena, Andy Garcia, Abdiel Vasallo Veliz, Sofia Leyva, Valentina Roldan, Robert P. Vertes, Timothy A. Allen

## Abstract

Using episodic memories to help guide decisions requires top-down medial prefrontal cortex interactions with the hippocampus. Mechanistically, this integrated prefrontal-hippocampal memory state is thought to be organized by synchronized network oscillations and mediated by connectivity with the nucleus reuniens of the thalamus. Here, we recorded local field potentials from the prefrontal-reuniens-hippocampal network while rats were engaged in a nonspatial sequence memory task which helps isolate memory-related activity from running-related oscillations. We found that synchronous prefrontal-hippocampal beta bursts (15-30 Hz) were dominant during memory trials, whereas synchronous theta (6-12 Hz) was highest during running. Beta rose during a trial and peaked just before a decision, whereas theta was highest during running. Beta bursts first appeared in reuniens and then in prefrontal and hippocampal sites simultaneously, suggesting beta could be directly driven by reuniens. To test this, we used an optogenetic approach (retroAAV-ChR2) to see if reuniens was capable of driving prefrontal-hippocampal beta synchrony. Reuniens activation induced prefrontal-hippocampal beta coherence, and reduced theta coherence, resembling the observed memory-driven network state. These findings demonstrate for the first time that reuniens contributes to memory by driving transient synchronous beta in the prefrontal-hippocampal system facilitating coherent interactions that sub-serve memory-based decision making.

## INTRODUCTION

Episodic memory is adaptive in its ability to advantage future behaviors, which depend on interactions between the medial prefrontal cortex and the hippocampus (Chiba, Kesner and Gibson, 1997; Allen and Fortin, 2013; Eichenbaum, 2017; Jayachandran *et al*., 2019; Reeders *et al*., 2021). A general axiom contends that prefrontal cortex exercises control over memories that are otherwise represented with their spatiotemporal contexts in the hippocampus (Ferbinteanu, Kennedy and Shapiro, 2006; Preston and Eichenbaum, 2013). That is, information relayed from the prefrontal cortex to the hippocampus guides situation-specific acquisition and retrieval, while in turn, the hippocampus provides context-specific memory in the form of episodes, goals, rules, and procedures (Ranganath and Ritchey, 2012; Allen and Fortin, 2013; MacDonald *et al*., 2014; Davachi and DuBrow, 2015; Ito *et al*., 2015). Theoretically, this involves using working memory to help actively buffer long-term memories so they can be suitably manipulated for use in decision making (Baddeley, 2000).

Anatomically, higher-order thalamocortical connections through the nucleus reuniens likely contribute to controlling these memory dynamics (Gazzaley and D’Esposito, 2007; Theyel, Llano and Sherman, 2010; Poulet *et al*., 2012; Ito *et al*., 2015; Viena, Linley and Vertes, 2018; Jayachandran *et al*., 2019; Malik *et al*., 2022). While, the hippocampus directly projects to the prefrontal cortex (Hoover and Vertes, 2007), only sparse return projections have been identified (Sesack *et al*., 1989; Laroche, Davis and Jay, 2000; Rajasethupathy *et al*., 2015; Malik *et al*., 2022). Instead, memory-related interactions depend on the nucleus reuniens, a region of the midline thalamus that provides a bi-directional control over prefrontal-hippocampal loops (Vertes *et al*., 2006; Varela *et al*., 2014; Dolleman-van der Weel *et al*., 2019; Viena, Vertes and Linley, 2021). Several labs have shown that reuniens is critical for regulating memory acquisition and retrieval (Xu and Südhof, 2013; Hallock, Wang and Griffin, 2016; Jayachandran *et al*., 2019), but how it supports memory is unclear. The most prevalent theory is that reuniens organizes oscillatory synchrony in order to mediate prefrontal-hippocampal interactions (Eichenbaum, 2017). Consistent with this notion, reuniens can impose slow oscillations in CA1, coupled to slow waves including delta (0.1-4 Hz; Zhang *et al*., 2012; Hauer, Pagliardini and Dickson, 2019; Angulo-Garcia *et al*., 2020), phase lock to hippocampal theta (6-12 Hz) during spatial alternations (Varela *et al*., 2014; Ito, Moser and Moser, 2018), and plays a critical role in organizing distal gamma bursts (30-90 Hz; (Ferraris *et al*., 2018). But how and when prefrontal-hippocampal networks synchronize during memory, and how this is related to reuniens, remains poorly understood.

Theta oscillations have been extensively studied in prefrontal-hippocampal-dependent memory research (Buzsáki, 2002; Hyman *et al*., 2005; Jones and Wilson, 2005; Kim *et al*., 2011; O’Neill, Gordon and Sigurdsson, 2013; Gonzalez *et al*., 2022) and have been shown to organize memory dynamics throughout the prefrontal-reuniens-hippocampal network (Siapas, Lubenov and Wilson, 2005; Colgin, 2011; Hyman, Hasselmo and Seamans, 2011; Varela *et al*., 2014; Roy *et al*., 2017; Griffin, 2021; Hauer, Pagliardini and Dickson, 2022). In general, theta coordinates brain activity during waking behaviors and in REM sleep across the phylogeny, and appears across physiological levels of analysis, suggesting it as a common means of organizing synaptic integration across distal locations in the brain (Raghavachari *et al*., 2001, 2001; Buzsáki, 2002; Buzsáki and Draguhn, 2004; Allen and Fortin, 2013; Hsieh and Ranganath, 2014). Paired prefrontal-hippocampal recordings from awake behaving rats have shown that prefrontal neurons are less hippocampal theta entrained when reuniens is inactivated (Hallock, Wang and Griffin, 2016), and this impairs spatial working memory. Reuniens has also been shown to have prominent theta during REM sleep (Viena, Vertes and Linley, 2021). Moreover, theta contributes to memory consolidation, spatial navigation, nonspatial sequence memory, and working memory (Buzsáki, 2002; Buzsáki and Moser, 2013; Zhang and Jacobs, 2015; Herweg, Solomon and Kahana, 2020; Shahbaba *et al*., 2022) - all tasks that require reuniens. However, our understanding of rhythmic prefrontal-reuniens-hippocampal synchrony may be limited due, in part, to the fact that most memory tasks use running behaviors that alone drive theta oscillations irrespective of memory (Hyman *et al*., 2005; Allen *et al*., 2016; Schultheiss *et al*., 2020; Kennedy *et al*., 2022).

Although less studied, there is also evidence that beta rhythms (15-30 Hz) synchronize hippocampal memory networks (Martin, Beshel and Kay, 2007; Igarashi *et al*., 2014; Allen *et al*., 2016; Symanski *et al*., 2022). In rats, beta is often seen in the hippocampus during odor-based memory tasks and has also been associated with long-term odor memory (Martin, Beshel and Kay, 2007) and temporal contexts (Allen *et al*., 2016). Beta organizes CA1-lateral entorhinal cortex interactions (Igarashi *et al*., 2014), is stronger in proximal CA1 late in a trial (Gattas *et al*., 2022), and entrains hippocampal spiking activity during memory trials that are otherwise theta entrained during running (Igarashi *et al*., 2014; Allen *et al*., 2016). Notably, beta is also found in the olfactory bulb but is lost there when centrifugal inputs are cut off (Martin, Beshel and Kay, 2007). Olfactory bulb-hippocampal coherence is not clearly related to memory and decision making but may develop in parallel (Symanski *et al*., 2022). Further, hippocampal beta is low early in learning and higher when memory is stable and engaged in a flexible manner (Martin, Beshel and Kay, 2007; Igarashi *et al*., 2014; Allen *et al*., 2016). The available data suggests there are central drivers orchestrating beta in support of memory-related functions, such as sensorimotor integration and/or working memory (Lundqvist *et al*., 2016; Miller *et al*., 2018; Schmidt, Duin and Redish, 2019). Along these lines, cortical beta may serve as a general mechanism for maintaining synaptic information states during stimulus-free memory delays, helping to bridge time and aid subsequent decisions (Lundqvist *et al*., 2016). Moreover, thalamocortical oscillations have been related to reducing sensory-sourced influences and shifting toward internal processing states in beta (Poulet *et al*., 2012). However, whether and how beta synchronizes in prefrontal-hippocampal regions, and the role of beta in reuniens, is not known.

Here, we tested if and when the prefrontal-reuniens-hippocampal network synchronize during nonspatial sequence memory focusing primarily on beta and theta. We used a nonspatial sequence memory task that models the ‘flow of events’ (episodic memory) to help separate memory-related activity from running/navigational activity (Allen *et al*., 2014, 2016; Jayachandran *et al*., 2019, 2022). We found that beta dominated the prefrontal-reuniens-hippocampal network during memory trials, whereas theta dominated only during running behaviors. The time course suggested reuniens could drive beta in the prefrontal-hippocampal network as it came on both earlier and stronger. Thus, we tested if reuniens could drive network synchrony directly using an optogenetic approach (retroAAV-ChR2). We found that optogenetic activation of reuniens (at any frequency) was sufficient to induce beta coherence and reduce theta coherence across the prefrontal-hippocampal system. The pattern of reuniens-induced prefrontal-hippocampal synchrony was remarkably similar to the network state observed during sequence memory trials, suggesting reuniens activity alone can trigger prefrontal-hippocampal beta memory states. In sum, these results suggest a model in which reuniens imposes transient beta bursts onto the prefrontal-hippocampal system helping coordinate a synchronous state that is conducive to using episodic-like sequence memories for decision-making.

**RESULTS**

Performance on the sequence task requires rapid sequential context retrievals from long-term memory. This type of sequence memory depends on the prefrontal-reuniens-hippocampal system (Jayachandran *et al*., 2019), and shows neurocognitive parallels between rats and humans (Allen *et al*., 2014; Reeders *et al*., 2021). The sequence task alternated between two sequence lists (ABCD and WXYZ), presented at opposite ends of a straight-alley maze to help eliminate the possibility that rats could hold a single sequence in mind throughout the session, further pressuring directed retrievals (see Methods).

We recorded from paired prefrontal-hippocampal and reuniens-hippocampal sites during the nonspatial sequence memory task to characterize rhythmic activity throughout the system. After asymptotic training on both sequences, rats were implanted with chronic silicon probes. The first group had dual prefrontal-hippocampal implants (*n* = 4), and the second group had dual reuniens-hippocampal implants (*n* = 4). One additional rat had a single implant in prefrontal cortex only and was included in analyses where dual-site implants were not critical. All implants were composed of two 32-channel silicon probes (NeuroNexus and Cambridge NeuroTech) that were positioned in deep layers of prelimbic cortex (prefrontal cortex, **Figure S1A**_**i**_), the nucleus reuniens of the thalamus (**Figure S1B**_**i**_), and/or stratum lacunosum moleculare layers in dorsal CA1 of the hippocampus (slm, **Figure S1A**_**ii**_**B**_**ii**_). Anatomically, these regions were chosen because they are directly interconnected through glutamatergic projections, with the exception that there are sparse prefrontal projections to CA1. Reuniens projections innervate both glutamatergic and GABAergic target cells in prefrontal cortex and the slm in CA1, and thus reuniens can control the excitatory-inhibitory tone in these regions (for review see Dolleman-van der Weel *et al*., 2019).

We focused on local field potentials (LFPs) here because they provide insight into rhythmic network dynamics occurring during memory and reflect synchronous barrages of synaptic field activity near the electrode. Analysis windows were primarily centered around high memory demand trials (positions 2-4) and sorted by sequential contexts and accuracy. We also examined running and stationary (awake) behaviors to compare low memory behaviors with behaviors during memory (odor trials).

During recordings, rats demonstrated sequence memory by making in-sequence (poke > 1 s; InSeq) and out-of-sequence (poke < 1 s; OutSeq) decisions in the nosepoke port one odor at a time (**Figure 1A**). Odors were terminated by the rat at withdrawal, or after 1 s. Position 1 was excluded from the memory analysis because items were never presented OutSeq in that position and had no memory demand beyond the rule that it was the start of the sequence list. By contrast, OutSeq odors were presented in positions 2-4, and thus these were considered high memory demand trials. Global sequence memory was measured using a sequence memory index (SMI) which weighs the relative response patterns on InSeq and OutSeq responses (see Allen *et al*., 2014; Jayachandran *et al*., 2022 for details) and provides a single value for sequence memory ranging from -1 to 1, where 1 represents perfect sequence memory. Using this measure, we found that sequence memory was high throughout the experiments in all rats (*n* = 9 rats; experiment-wide SMI = 0.28 ± 0.01; t_(26)_ = 24.62, *p* = 1.53×10^−19;^ all individual subject G-tests < 0.05), at levels expected based on previous studies.

**Figure 1.**
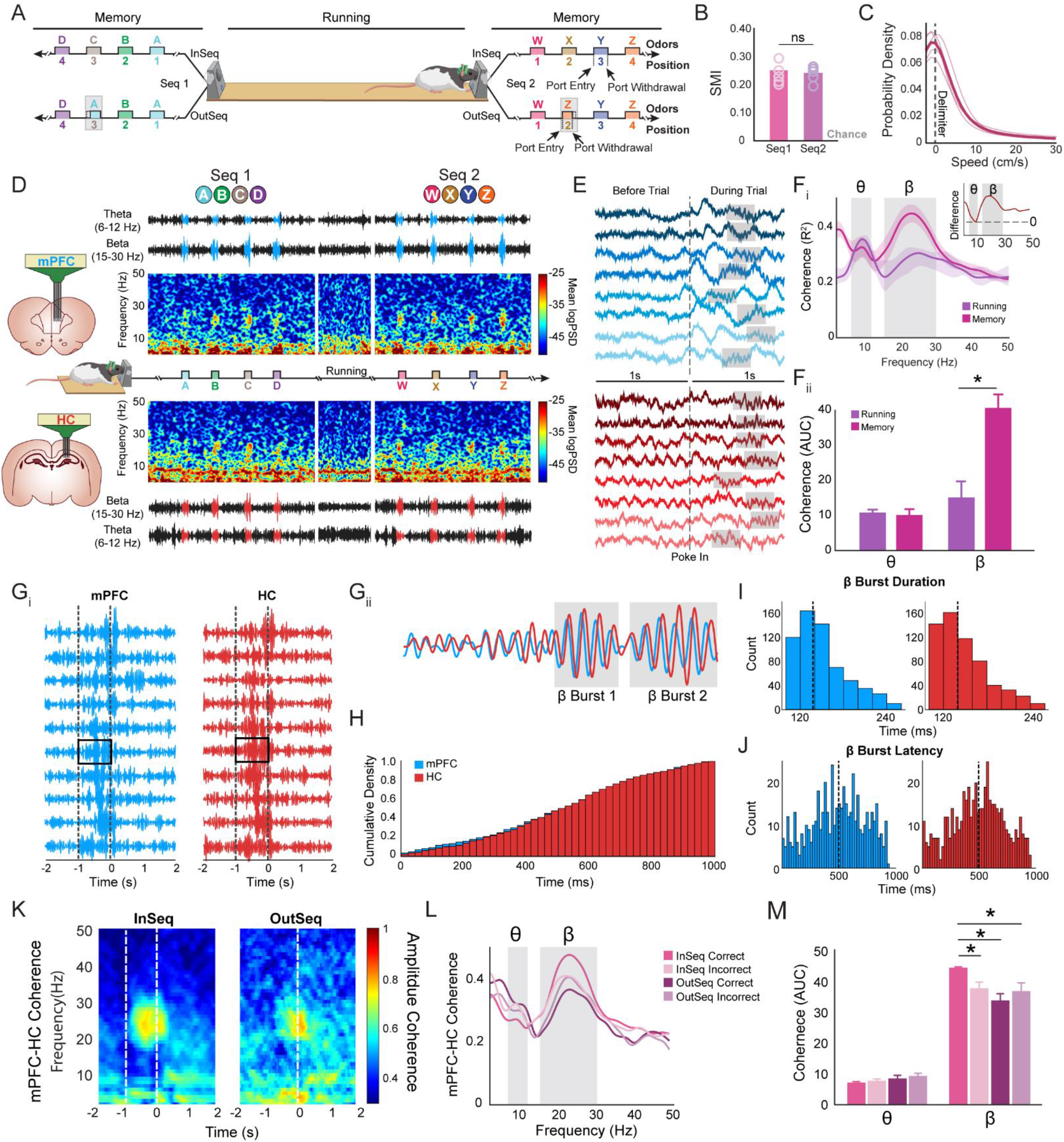
Prefrontal-hippocampal system shows transient beta coherence in nonspatial sequence memory. **A)** A linear track was used with odor ports located at opposite ends where two separate four-odor sequences (A, B, C, D or W, X, Y, Z) were presented. Rats had to correctly identify the odor as either InSeq (hold ≥ 1 s) or OutSeq (hold < 1 s). **B)** SMI was not significantly different between Seq 1 and Seq 2. Individual rat performance is indicated by circles. **C)** Average running speeds less than 10 cm/s represent 80% of the data. **D)** Representative spectrogram with corresponding filtered LFP from theta and beta bands in prefrontal cortex (mPFC, blue) and hippocampus (HC, red) during both sequences interleaved with a short running bout. High power beta is aligned to the odor sampling period. **E)** Sample raw LFP (two samples from each rat within the experiment) in prefrontal cortex (blue) and hippocampus (red). Each rat is indicated with a different shade of color. Beta bursts are highlighted in gray and can be seen only during the trial (after poke in). **F**_**i**_) Prefrontal-hippocampal coherence was significantly different between running periods (non-memory) and memory (odor-trials). Inset shows a difference between memory and running. Beta synchrony increases during memory while theta was lower (close to zero). **F**_**ii**_) AUC shows that theta coherence was not significantly different between running and memory, while beta coherence was significantly different between running and memory. **G**_**i**_) Sample bandpass beta filtered from prefrontal and hippocampal sites with a sample of 10 trials shows closely matched high amplitude beta ∼100ms after the poke in. **G**_**ii**_) A zoomed in trial shows the bursty properties of beta. **H)** The probability of prefrontal and hippocampal beta burst occurrence are not significantly different. **I)** Average duration of a beta burst was not significantly different between prefrontal and hippocampal sites. **J)** The latency to the first beta burst was not significantly different between prefrontal and hippocampal sites. **K)** Prefrontal-hippocampal coherence separated based on sequential context (InSeq vs OutSeq). **L)** Coherence between InSeq and OutSeq trials separated based on accuracy. **M)** Beta AUC was significantly different across the four trail types. InSeq_correct_ trials was significantly higher compared to InSeq_incorrect_, OutSeq_correct_, and OutSeq_incorrect_. Theta AUC was not significantly different across the four trial types. Abbreviations: mPFC, medial prefrontal cortex; HC, hippocampus, InSeq, in-sequence; OutSeq, out-of-sequence; SMI, sequence memory index; Seq, sequence; LFP, local field potential; AUC, area under the curve; InSeq_correct_, in-sequence correct, InSeq_incorrect_, in-sequence incorrect, OutSeq_correct_, out-of-sequence correct, OutSeq_incorrect_, out-of-sequence incorrect; θ, theta; β, beta; ns, not significant.

### Sequence memory and running behaviors

In all experiments, recordings began after sequence memory performance reached pre-surgical levels (t_(74)_ = -1.16, *p* = 0.25). During recording sessions, sequence memory was evident across all positions (Pos2: 0.43±0.03; Pos3: 0.35±0.04; Pos4: 0.32 0.06) and odors (OdorA: 0.43±0.05; OdorB: 0.39±0.06; OdorC: 0.34±0.08; OdorD: 0.35±0.08; OdorW: 0.50±0.04; OdorX: 0.35±0.12; OdorY: 0.46±0.06; OdorZ: 0.38±0.08). Performance was similar on sequence 1 (0.28±0.03) and sequence 2 (0.25±0.02), with no significant difference between them (t_(53)_= -0.80, *p* = 0.43), highlighting the rats ability to remember both sequences, and ability to switch between sequences. In the prefrontal-hippocampal rats there was no significant difference in performance between sequence 1 and sequence 2 (t_(4)_ = 0.22, *p* = 0.84, **Figure 1B**). Furthermore, non-mnemonic aspects of behavior did not differ across sessions **(Figure S2A**).

Because it has been shown that distinct rhythmic network modes exist in the prefrontal-hippocampal circuit distinguished by animals’ gross behavioral states (e.g., Schultheiss *et al*., 2020), we divided our analyses into three behavioral states: (1) memory (sequence trials), (2) running, and (3) stationary periods using an open-source machine learning tracking algorithm (DeepLabCut). Running was set as speeds > 2.5 cm/s, and stationary bouts were set as < 2.5 cm/s, limited to when the animal was located on the linear track rather than in the nosepoke area. In this task, we found the probability density function of speeds (cm/s) < 10 cm/s accounted for ∼80% of the data (**Figure 1C**). See **Figure S3A** for sample tracking and behavioral segregations.

### Beta increases in the prefrontal cortex and hippocampus during sequence memory trials

We next focused on low frequencies (1-50 Hz) and on *a priori* defined beta (15-30 Hz) and theta (6-12 Hz) bands, given a previous report of their relevance in the sequence task in the hippocampus (Allen *et al*., 2016). **Figure 1D** shows representative spectrograms from the prefrontal cortex and hippocampus during sequence 1 and sequence 2 interleaved by a short representative running sample. Overall, prefrontal sites were dominated by power in delta and theta. However, upon initiating a trial, we observed brief intermittent beta clouds (∼15-30 Hz). Hippocampal sites were similar with the exception that more power was seen in theta and less in delta, but again beta clouds were aligned with prefrontal cortex and occurred only during a trial. This pattern repeated itself during the sequence task, with spectrograms rapidly transitioning from theta dominating during running to beta dominating during sequence trials. The magnitude of the transient beta activity in both prefrontal cortex and hippocampus was high, as seen in the large signal-to-noise ratio in the beta-filtered voltage traces (**Figure 1D**). Compared to theta, beta amplitudes increased drastically on the sequence memory trials.

We looked at the raw voltage traces of prefrontal cortex and hippocampus and saw the same rhythmic patterns providing further compelling evidence that beta bursts occur in both the prefrontal cortex and hippocampus during odor sampling in the sequence task (**Figure 1E**). Upon trial initiation the prefrontal cortex developed large delta waves accompanied by brief intermittent beta bursts. In the hippocampus, theta oscillations can be seen occurring before trial initiation and appear to continue for ∼3 cycles, but the traces are then interposed with intermittent beta bursts. Notably, the prefrontal and hippocampal paired raw traces appear to show a rough trial-by-trial alignment suggesting long-distance synchrony.

We performed a coherence analysis to further evidence beta synchrony between prefrontal-hippocampal electrode pairs, sorting high memory demand odor trials (positions 2-4) and running periods (**Figure 1F_i_**). First, aggregate prefrontal-hippocampal coherence values were calculated and plotted across frequencies in 0.4883 Hz steps. We found the peak coherence frequencies, on average, were at 22.34±0.81 Hz (individual peaks = 23.92, 23.43, 20.51, and 21.48 Hz) during memory trials, and 10.01±0.65 Hz (individual peaks = 10.25, 10.74, 9.77, 8.79 Hz) during running. The magnitudes of these peaks were significantly higher during memory trials compared to running periods (t_(3)_ = 13.71, *p* = 9.00 × 10^−6^). Next, we looked at the maximum coherence magnitudes (R^2^) found in the beta band during memory and compared them with those found in the theta band during running as a benchmark for the strength of the synchrony. Beta coherence magnitudes during odor trials (memory-related β_coherence_ = 0.37±0.021), were comparable to theta values during running (running-related θ_coherence_ = 0.35±0.01; t_(3)_ = 1.30, *p* = 0.24). Next, we ran a behavior x frequency two-way ANOVA to see if the overall magnitude or shape of the coherence plots were different in memory and running periods. There was a significant effect of the behavioral condition (F_(1,824)_ = 660.42, *p* = 1.27×10^−99^) with higher overall coherence values during memory compared to running conditions, a main effect of frequency (F_(102,824)_ = 6.36, *p* = 8.05×10^−50^) which partially reflect the 1/f noise, and a significant behavior x frequency interaction effect (F_(102,824)_ = 3.38, *p* = 1.31×10^−20^). Thus, prefrontal-hippocampal synchrony showed an overall increase during memory trials as well as showed that separate frequencies dominate during memory and running. This analysis provides strong support for the notion that prefrontal-hippocampal synchrony shifts between brief beta states during memory, and theta states during running. Furthermore, we plotted a difference function (memory vs. running) to identify the specific frequencies that synchronize or desynchronize (**Figure 1F_i_**, inset) revealing that beta synchrony increases during memory (t_(3)_ = 21.05, *p* = 2.35×10^−4^), while theta was trending lower although this did not reach significance (t_(3)_ = -2.99, *p* = 0.06). Notably, the frequency shifts were very closely matched to our pre-defined frequency bands for beta (15-30 Hz) and theta (6-12 Hz). Lastly, we analyzed beta and theta as a whole band using area under the curve (AUC) measurements (**Figure 1F_ii_**). This also showed that beta increased during memory (t_(6)_ = 4.03, *p* = 0.01), the theta was not significantly different between conditions (t_(6)_ = -0.47, *p* = 0.65). We then tested the possibility that beta coherence might simply reflect the fact that the animal is stationary during memory trials. We demonstrate that memory trials had significantly more beta coherence compared to other stationary periods (not engaged in the trials) during the task (**Figure S4A**). Together, these results show that the prefrontal-hippocampal synchrony during sequence memory is predominantly driven by transient increases in beta.

### Prefrontal-hippocampal beta bursts occur simultaneously and late in a memory trial

We sought to further describe the co-occurring beta in both prefrontal cortex and hippocampus during odor trials in the sequence memory task. First, we plotted bandpass beta filtered traces from prefrontal and hippocampal sites on a trial-by-trial basis to examine beta in both regions (sample in **Figure 1G_i_**, also see **Figure S5** for all animals). On every trial, the beta traces in both prefrontal cortex and hippocampus were very closely matched in both time and amplitude. High amplitude beta consistently appeared later (∼100 ms) in a trial. We then overlaid prefrontal cortex and hippocampal traces and looked at cycle-by-cycle changes in the beta amplitudes and phases between regions. When amplitudes were small (in between trials or early in the 1 s odor trials), there was modest to no phase or amplitude synchrony. When amplitudes were large, prefrontal cortex and hippocampus were highly phase and amplitude synchronous (**Figure 1G_ii_**).

These beta cycles were bursty in nature rather than continuous, thus we developed a beta burst detection algorithm (custom-written scripts in MATLAB, see methods). We plotted the cumulative density function (CDF) of beta bursts during the sequence trials in the prefrontal cortex and overlaid it with the CDF from hippocampus. The CDF in prefrontal cortex and the hippocampus were almost identical (**Figure 1H**, t_(609)_ = 0.020, *p* = 0.98). Additionally, the average duration of a beta burst in prefrontal cortex and hippocampus were not significantly different (prefrontal: 147.95±2.29 ms hippocampus: 149.70±2.07 ms; t_(1239)_ = -0.71, *p* = 0.48; **Figure 1I**). The latency to the first burst onset upon trial initiation in prefrontal cortex was also not significantly different from hippocampus (prefrontal: 463.13±10.08 ms; hippocampus; 464.08±11.73 ms; t_(1022)_= -0.06, *p* = 0.95; **Figure 1J**).

### Prefrontal-hippocampal beta coherence magnitudes reflect temporal context memory

The long latencies of the bursts suggested that prefrontal-hippocampal beta could be involved in the mnemonic aspects of the sequence task. Thus, we analyzed prefrontal-hippocampal beta coherence as a function of the sequential context of the odors (InSeq or OutSeq). Perievent coherograms were aligned to nosepoke withdrawals and averaged over all InSeq and OutSeq trials ± 2 s (**Figure 1K**). InSeq trials show a large magnitude beta cloud (∼20-30 Hz) with a centroid in the second half of an odor trial as expected. On OutSeq trials pre-withdrawal beta is weaker and shorter. While this is consistent with prefrontal-hippocampal beta coherence differentiating the sequence context of odors this may simply be due to differences in the poke times (> 1 s for InSeq odors, and < 1 s for OutSeq odors). Thus, we separated by both sequential context and accuracy resulting in four conditions (**Figure 1L**). While each condition shows very high coherence (R^2^ > 0.3), InSeq_correct_ trials were consistently large in magnitude (similar to hippocampal amplitudes described in Allen et al., 2016). We then calculated AUC values within *a priori* beta and theta bands (**Figure 1M)** and tested for differences across conditions. We found that beta was significantly different across the four trial types (F_(3,15)_ = 4.24, *p* = 0.03). Post hoc analysis revealed InSeq_correct_ trials were significantly higher compared to InSeq_incorrect_ (*p* = 0.02), OutSeq_correct_ (*p* = 0.01), and OutSeq_incorrect_ (*p* = 0.02) trials. To control for poke time differences, we compared OutSeq_incorrect_ trials to InSeq_correct_ trials which have the same poke times (>1 s) and found a significant difference (*p* = 0.016). If longer poke times caused increased beta coherence, we would expect to have more beta during OutSeq_incorrect_ trials. Further, the OutSeq_correct_ (< 1 s) trials were not significantly different from OutSeq_incorrect_ (> 1 s) trials (*p* = 0.72). We found that theta was not significantly different across any of the four trial types (F_(3,15)_ = 1.74, *p* = 0.21). To further support the notion that beta is related to sequence memory, we ran an average perievent power spectral densities sorted by sequential contexts and accuracy for prefrontal cortex and hippocampus separately, all of which show evidence that local beta is related to sequence memory as well (**Figure S6A**). Observation of these prefrontal-hippocampal beta coherence suggests beta in the hippocampus may be associated with processing the temporal context of items, and, for the first time, shows that beta is engaged in prefrontal-hippocampal synchrony. However, the question remained what could be driving prefrontal-hippocampal beta synchrony?

### Beta increases in the nucleus reuniens of the thalamus during sequence memory trials

We hypothesized the nucleus reuniens of the thalamus could drive synchronous beta bursts in the prefrontal cortex and hippocampus. Reuniens has the necessary anatomy, including a population of dual prefrontal-hippocampal projecting cells, in order to drive this synchrony (Hoover and Vertes, 2007; Varela *et al*., 2016; Viena *et al*., 2021). Additionally, reuniens has been shown to be critical to sequence memory (Jayachandran *et al*., 2019). Therefore, we next performed dual-site reuniens-hippocampal recording during the sequence memory task.

In reuniens-hippocampal rats, recordings began after driving down probes to their target locations and after sequence memory performance matched pre-surgical levels, during which rats had strong and steady memory on both sequences (t_(3)_ = -0.03, *p* = 0.98; **Figure 2A**). We ran parallel behavioral analyses in reuniens-hippocampal rats to those ran in prefrontal-hippocampal rats for direct comparison (**Figure S2B**). We then looked at the overall spectrograms. **Figure 2B** shows a representative spectrogram from the reuniens during sequence 1 and sequence 2, interweaved by a short representative running sample. Note the hippocampus looked the same in these rats and thus these spectrograms are not shown here. In reuniens, the spectrograms were dominated by sporadic periods of high delta and theta frequencies that sometimes overlapped in time, and beta activity. Reuniens beta was broader (∼12-30 Hz) than the activity observed on hippocampal and prefrontal sites, but similarly to these regions, reuniens beta appeared repeatedly during trials throughout the sequence task recordings. The filtered voltage traces of reuniens showed the highest beta amplitudes are time-locked to trials of the sequence task (**Figure 2B**).

**Figure 2.**
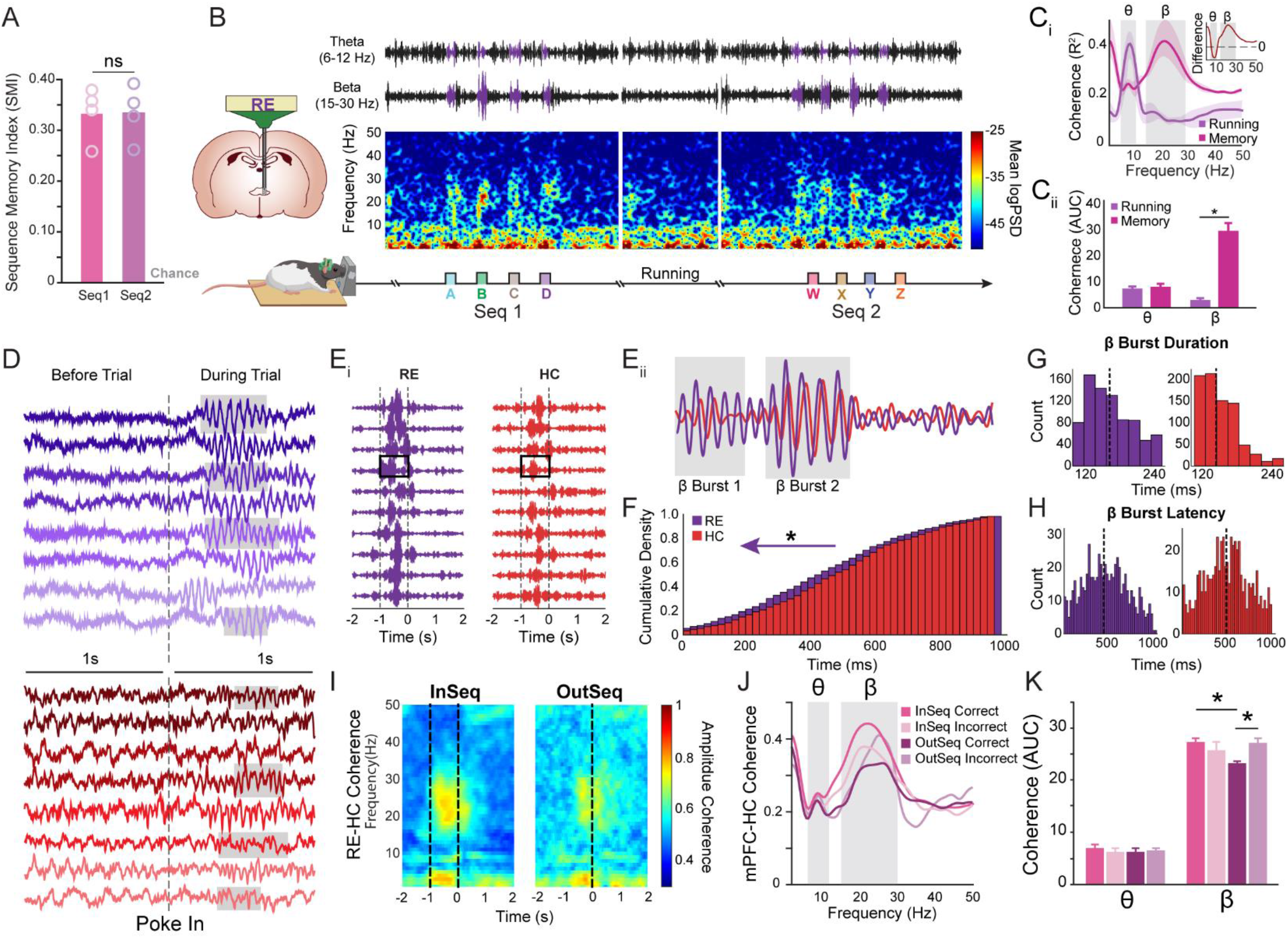
Nonspatial sequence memory induces beta bursts in reuniens. **A)** SMI was not significantly different between Seq 1 and Seq 2. Individual rat performance is indicated by circles. **B)** Representative spectrogram with corresponding filtered LFP from theta and beta bands in reuniens (RE; purple) during both sequences with a sample running bout in between. High power beta is aligned to the odor sampling period. **C**_**i**_) Reuniens-hippocampal coherence was significantly different between running periods (non-memory) and memory (odor-trials). Inset shows a difference (memory – running) and show that beta synchrony increases during memory while theta was lower (close to zero). **C**_**ii**_) AUC shows that theta coherence was not significantly different between running and memory, while beta coherence was significantly different between running and memory. **D)** Sample raw LFP (two samples from each rat within the experiment) in reuniens (purple) and hippocampus (red). Each rat is indicated with a different shade of color. Beta bursts are highlighted in gray and can be seen only during the trial (after poke in). **E**_**i**_) Sample bandpass beta filtered from reuniens and hippocampal sites with a sample of 10 trials shows reuniens beta occurs earlier than hippocampus beta. **E**_**ii**_) A zoomed in trial shows the bursty properties of beta with reuniens beta occurring earlier in the trial. **F)** The probability of reuniens and hippocampal beta burst occurrence are significantly different with reuniens occurring earlier indicated with purple arrow. **G)** Average duration of a beta burst was significantly different longer in reuniens than hippocampal sites. **H)** The latency to the first beta burst was significantly earlier in reuniens than hippocampal sites. **I)** Reuniens-hippocampal coherence separated based on sequential context (InSeq vs OutSeq). **J)** Coherence between InSeq and OutSeq trials separated based on accuracy. **K)** Beta AUC was significantly different across the four trail types. InSeq_correct_ trials were significantly higher compared to OutSeq_correct_ and OutSeq_correct_ trials were significantly lower than InSeq_incorrect_. Theta AUC was not significantly different across the four trial types. Abbreviations: RE, nucleus reuniens; HC, hippocampus, SMI, sequence memory index; Seq, sequence; LFP, local field potential; AUC, area under the curve; InSeq, in-sequence; OutSeq, out-of-sequence; InSeq_correct_, in-sequence correct, InSeq_incorrect_, in-sequence incorrect, OutSeq_correct_, out-of-sequence correct, OutSeq_incorrect_, out-of-sequence incorrect; θ, theta; β, beta; ns, not significant.

We next performed a coherence analysis on reuniens-hippocampal electrode pairs, sorting high memory demand odor trials (positions 2-4) and running periods (**Figure 2C_i_**). First, aggregate reuniens-hippocampal coherence values were calculated and plotted across frequencies. We found the peak coherence frequencies, on average, were at 22.09±0.54 Hz (individual peaks = 23.44, 22.46, 20.99, and 21.48 Hz) during memory trials, and 8.26±0.21 Hz (individual peaks = 8.30, 7.79, 8.79, 8.19 Hz) during running. The magnitudes of these peaks were significantly higher during memory trials compared to running periods (t_(3)_ = 23.85, *p* = 3.57×10^−7^). Beta coherence magnitudes during odor trials (memory-related β_coherence_ = 0.42±0.08), were comparable to theta values during running (running-related θ_coherence_ = 0.40±0.06; t_(3)_ = 0.14, *p* = 0.89). Next, we ran a behavior x frequency two-way ANOVA to see if the overall magnitude or shape of the coherence plots were different in memory and running periods. There was a significant effect of the behavioral condition (F_(1,824)_ = 448.92, *p* = 2.61×10^−75^) with higher overall coherence values during memory compared to running conditions, a main effect of frequency (F_(102,824)_ = 4.28, *p* = 8.23×10^−30^) which partially reflect the 1/f noise, and a significant behavior x frequency interaction effect (F_(102,824)_ = 9.28, *p* = 1.61×10^−74^). Thus, reuniens-hippocampus reflects an increase in overall synchrony, driven by different frequencies bands during memory trials compared to running similar to prefrontal-hippocampal synchrony. This analysis provides strong support for the notion that reuniens-hippocampal synchrony also shifts between brief beta states during memory and theta states during running. We then plotted a difference function (memory vs. running) to identify the specific frequencies that synchronize or desynchronize (**inset Figure 2C_i_**). This analysis showed that beta synchrony increases during memory (t_(3)_ = 3.86, *p* = 0.03), while theta was trending lower however not significance (t_(3)_ = -2.12, *p* = 0.13). Lastly, we analyzed beta and theta as a whole band using AUC measurements (**Figure 2C_ii_**). This also showed that beta increased during memory (t_(6)_ = 9.48, *p* = 7.8×10^−5^), but theta was not significantly different between conditions (t_(6)_ = -1.79, *p* = 0.12), demonstrating that memory trials had significantly more beta coherence than in running. We also tested the possibility that beta coherence might simply reflect the fact that the animal is stationary during memory trials. However, we demonstrate that memory trials had significantly more beta coherence compared to other stationary periods during the task (**Figure S4B**).

We next looked at the raw voltage traces of reuniens and hippocampus showing modest delta and theta activity prior to trial initiation, followed by large beta bursts in reuniens after trial initiation (**Figure 2D**). Unlike the beta seen in hippocampal and prefrontal sites, reuniens beta appears stronger and more oscillatory in nature. Additionally, beta in reuniens appeared to have slightly earlier onset latencies than in prefrontal cortex and hippocampus.

### Reuniens beta bursts occur shortly before hippocampal beta bursts

If reuniens were driving prefrontal-hippocampal beta synchrony, the bursts in reuniens should occur *before* beta bursts in the hippocampus. To test the temporal relationships, we plotted the bandpass beta filtered traces in each structure and placed them next to each other on a trial-by-trial basis (sample in **Figure 2E_i_**, also see **Figure S5**). On each trial, high amplitude beta bursts in reuniens were easily identified and much larger than those observed in the hippocampus, and often occurred just before those in the hippocampal traces. We overlaid traces and zoomed in to look at the cycle-by-cycle changes in amplitude and phase comparing the two regions directly (**Figure 2E_ii_**). Early in the trial (e.g., beta burst 1 in **Figure 2E_ii_**) the phase and amplitudes were only modestly related. Later in the trial (e.g., beta burst 2 in **Figure 2E_ii_**) amplitudes were more tightly coupled.

Next, we quantified beta burst durations and latencies in both the reuniens and the hippocampus. The overlaid CDF of the beta bursts in each region shows reuniens bursts occur significantly earlier in a trial than in the hippocampus (t_(859)_ = 3.62, *p* = 3.17×10^−4^; **Figure 2F**). Additionally, we found the average duration of a beta burst was significantly longer in reuniens (177.08±3.25 ms) compared to the hippocampus (147.34±4.76 ms) (t_(1247)_ = 10.30, *p* = 5.98×10^−24^; **Figure 2G**), while the beta burst latency in reuniens was significantly earlier from hippocampus (reuniens: 393.00±8.23 ms; hippocampus; 431.50±10.60 ms; t_(1231)_ = -3.137, *p* = 0.002; **Figure 2H**).

### Reuniens-hippocampal beta coherence differs across sequence task conditions

We then analyzed reuniens-hippocampal coherence as a function of the sequential context. The coherograms showed strong beta on InSeq compared to OutSeq trials (**Figure 2I**). We then plotted the aggregate coherence sorted by sequential context and accuracy (**Figure 2J**). While theta was not significantly different across the four trial types (F_(3,15)_ = 0.27, *p* = 0.84), beta was significantly different (**Figure 2K**; F_(3,15)_ = 4.59, *p* = 0.02). Post hoc analysis showed that InSeq_correct_ trials were significantly more beta coherent compared to OutSeq_correct_ trials (*p* = 0.01). Additionally, OutSeq_correct_ trials were significantly lower than OutSeq_incorrect_ trials (*p* = 0.01). There were no significant differences between InSeq_correct_ and OutSeq_incorrect_ trials, which have the same hold time (*p* = 0.97). Lastly, InSeq_correct_ trials were not significantly different from InSeq_incorrect_ trials (*p* = 0.27). Thus, while there were differences in reuniens-hippocampal beta coherence, the fact that beta was consistently higher on the longer poke requirements are inconclusive about whether this coherence differentiates mnemonic content. Adding to this, **Figure S6B** shows the perievent spectrograms for each brain region individually, with reuniens being the only site with significant beta power on OutSeq incorrect trials, suggesting reuniens activity may be task- and hippocampal-related. Thus, reuniens may potentially engage prefrontal-hippocampal memory networks but itself be more driven by the behavioral dynamics inherent in the task (such as the poke and hold behavior).

### Time course of beta and theta activity in the prefrontal cortex, reuniens, and hippocampus in sequence memory trials

We wanted to directly compare the relative amplitudes and time courses for beta and theta across the three brain regions, providing a network wide perspective (**Figure 3A**). We took the upper envelope from the beta and theta filtered voltage traces, and *z*-normalized the output. **Figure 3A** shows an example upper envelope calculation for each band. These envelopes were then averaged across trials (InSeq_correct_ only to match poke times), and reflect a combination of the probability densities of bursts and changes in amplitudes. In reuniens, beta amplitudes start rising, on average, just prior to the onset of a trial and before prefrontal cortex and the hippocampus. Prefrontal cortex and hippocampus beta activity rises only after the trial starts but does so simultaneously (**Figure 3B**). We first compared the hippocampal recordings from the two experiments and found no significant difference between region but significantly different across time (region: F_(1,3)_ = 0.28, *p* = 0.63; time: F_(19,57)_ = 13.91, *p* = 5.61×10^−15^; region x time: F_(19,57)_ = 0.36, *p* = 0.99; inset **Figure 3B**). Therefore we combined all hippocampal rats and ran a brain region x time ANOVA showing a main effect of brain region (F_(2,16)_ = 33.24, *p* = 1.00×10^−3^), a main effect of time (F_(19,57)_ = 28.91, *p* = 1.71×10^−22^), and brain region x time interaction effect (F_(38,114)_ = 2.443, *p* = 1.48×10^−4^). This pattern indicates that (1) poking behaviors initiate beta in reuniens, and (2) reuniens drives prefrontal-hippocampal beta coherence. To provide further evidence, we calculated the slopes in 100 ms bins and tested for significantly increasing beta activity before, during, and after a trial (**Figure 3C**). In agreement, this analysis shows reuniens beta rising significantly before the start of a trial, where beta was decreasing before the start of a trial in prefrontal cortex and hippocampus, and significantly rising 100-600 ms into a trial (one sample t-test, *p* < 0.05). Interestingly all three brain regions appear to peak at the same time ∼500 ms and come down slightly or remain steady (near zero) for the remainder of the trial. Beta comes down in all brain regions to baseline levels starting about 100 ms after the trial ends.

**Figure 3.**
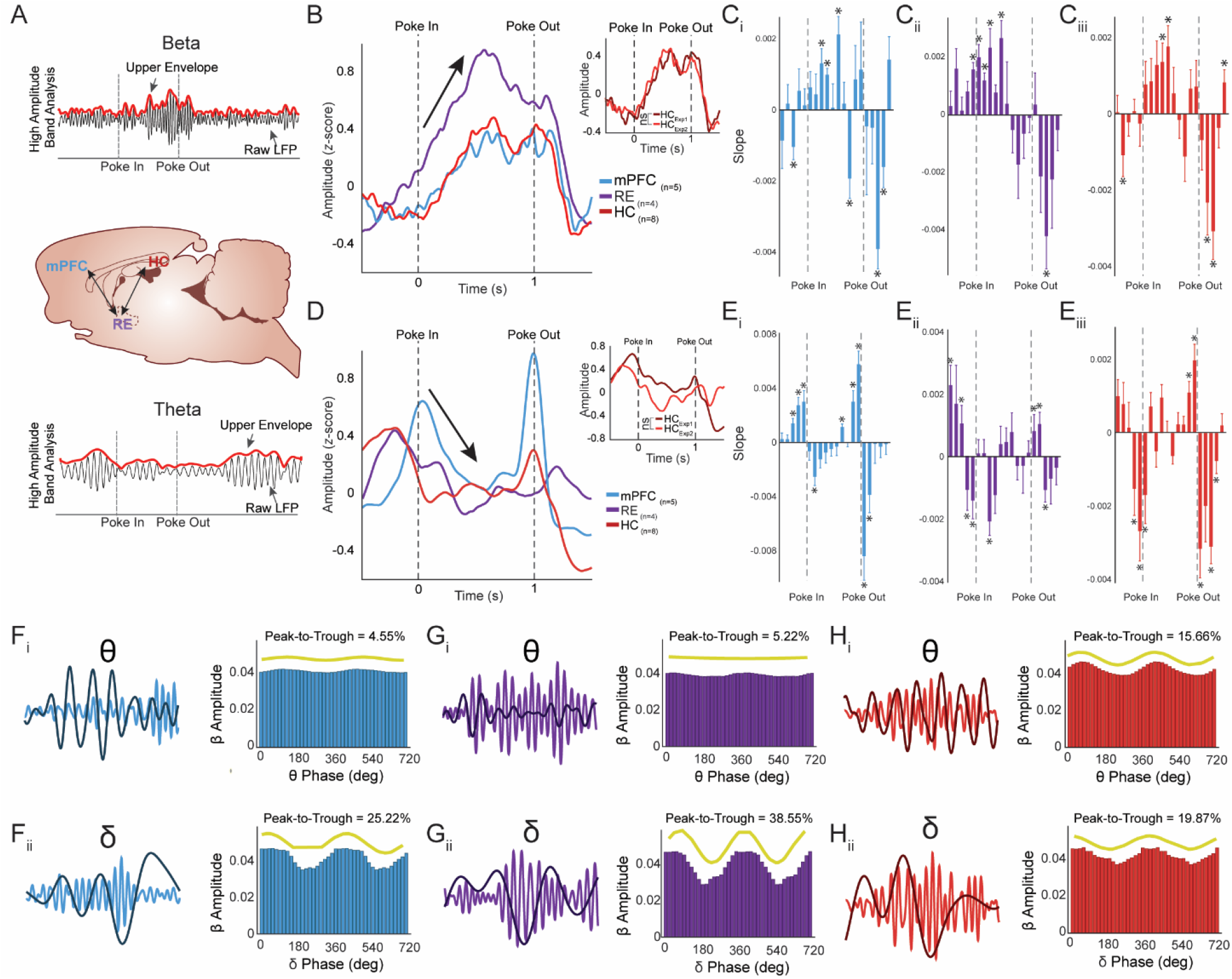
Beta amplitudes rise earlier and stronger in reuniens and then concurrently in prefrontal cortex and hippocampus. **A)** A sample of the upper envelope for both beta and theta for all three regions (prefrontal cortex, reuniens, and hippocampus). **B)** Beta amplitudes in reuniens start rising just prior to the onset of trials indicated with an arrow. Prefrontal cortex and hippocampus beta simultaneously rise after the trial. Overall showing a main effect of region, time and region x time. Inset plot shows that the hippocampal recordings from both data sets were not significantly different. **C**_**i**_) Calculated slopes show that in prefrontal cortex beta significantly rose ∼300 ms after the trial and then decreased after the poke out (*p*’s < 0.01, indicated with a star). **C**_**ii**_) In reuniens beta slopes significantly rose ∼100 ms before the onset of the trial and continued to rise during the trial (*p*’s < 0.01, indicated with a star). **C**_**iii**_) In hippocampus beta slopes shows similar patterns to prefrontal cortex, rising ∼300 ms after the trial (*p*’s < 0.01, indicated with a star). **D)** Theta amplitudes in reuniens and hippocampus show phasic increases before a trial followed by a phasic increase in prefrontal cortex at the start of the trial. During the trial there is a decrease in theta indicated with an arrow. Overall showing a main effect of region, time and region x time. Inset plot shows that the hippocampal recordings from both data sets were not significantly different. **E**_**i**_) Calculated slopes show that in prefrontal cortex theta significantly rose ∼300 ms before the start of the trial and ∼200 ms before the poke out (end of trial), while decreasing during the trial (*p*’s < 0.01, indicated with a star). **E**_**ii**_) In reuniens theta slopes significantly rose ∼500 ms before the onset of the trial and decreased during the trial (*p*’s < 0.01, indicated with a star). **E**_**iii**_) In hippocampus theta slopes shows similar patterns to reuniens and prefrontal cortex where theta slopes decrease during the trial (*p*’s < 0.01, indicated with a star). **F**_**i**_) A sample of prefrontal cortex beta voltage (blue) superimposed with theta voltage (dark blue) shows that beta and theta are weakly coupled. Example phase-amplitude plot from a rat showing beta amplitude was weakly modulated by theta phase. **F**_**ii**_) A sample of prefrontal cortex beta voltage (blue) superimposed with delta voltage (dark blue) shows that beta and delta are strongly coupled. Example phase-amplitude plot from a rat showing beta amplitude was more modulated by delta phase. **G**_**i**_) A sample of reuniens beta voltage (purple) superimposed with theta voltage (dark purple) shows that beta and theta are weakly coupled. Example phase-amplitude plot from a rat showing beta amplitude was weakly modulated by theta phase. **G**_**ii**_) A sample of reuniens beta voltage (purple) superimposed with delta voltage (dark purple) shows that beta and delta are strongly coupled. Example phase-amplitude plot from a rat showing beta amplitude was more modulated by delta phase. **H**_**i**_) A sample of hippocampal beta voltage (red) superimposed with theta voltage (dark red) shows that beta and theta are coupled. Example phase-amplitude plot from a rat showing beta amplitude was modulated by theta phase. **H**_**ii**_) A sample of hippocampal beta voltage (red) superimposed with delta voltage (dark red) shows that beta and delta are coupled. Example phase-amplitude plot from a rat showing beta amplitude was modulated by delta phase. Yellow waveforms represent the sine waves fitted to the amplitude– phase distributions. Abbreviations: mPFC, medial prefrontal cortex; RE, nucleus reuniens; HC, hippocampus; Exp, experiment; θ, theta; δ, delta; β, beta.

The time course of theta was quite different from beta, and in some ways the inverse. Theta showed brief phasic increases in amplitudes in reuniens and hippocampus before a trial, followed by a phasic increase in prefrontal around the start of the trial (**Figure 3D**). Theta decreases in all regions in the first half of a trial and stays near zero until the trial ends which is marked by a phasic increase in theta in the prefrontal cortex and hippocampus followed by reuniens. We compared the hippocampal recordings from the two experiments and found no significant difference between regions but significantly different across time (region: F_(1,3)_ = 3.19, *p* = 0.17; time: F_(19,57)_ = 7.52, *p* = 1.37×10^−9^; region x time: F_(19,57)_ = 1.69, *p* = 0.06; inset **Figure 3D**). We ran a brain region x time ANOVA showing a main effect of brain region (F_(2,6)_ = 0.08, *p* = 0.92), a main effect of time (F_(19,57)_ = 3.81, *p* = 4.60×10^−5^), and brain region x time interaction effect (F_(38,114)_ = 2.51, *p* = 9.30×10^−5^). We next calculated the slope in 100 ms bins and tested for significantly increasing theta activity before, during, and after a trial (**Figure 3E**). In agreement, this analysis shows prefrontal, reuniens, and hippocampal theta rising significantly before the start of a trial and around the poke out (one sample t-test, *p* < 0.05). Interestingly all three brain regions appear to peak and quickly decrease (around the poke in) and remain steady (near zero) during the trial. Theta then comes down peaks in all brain regions to around the end of the trial. This suggests theta is having the most influence in this network around trials, including the onset and offset, rather than during the trial.

To directly test for beta-theta relationships, we ran a phase-amplitude coupling analysis during trials (**Figure 3F-H**). We estimated the instantaneous phase during memory trials in theta and beta and plotted the beta envelope (amplitude) as a function of theta phase (**Figure 3F_i_, G_i_, H_i_**). We then fitted the plots to sine waves and calculated significant fits (p < 0.05). Overall this showed beta amplitudes were cross-frequency coupled to theta, as has been shown before in CA1 (Igarashi *et al*., 2014), but consistently weaker than coupling to delta (**Figure 3F_ii_, G_ii_, H_ii_**).

We calculated the peak-to-trough distance by subtracting the maximum beta amplitude (peak) by the minimum beta amplitude (trough). Then we calculated the percentage of the peak-to-trough distance and ran a brain region x frequency ANOVA showing a main effect of frequency (F_(1,34)_ = 32.50, *p* = 4.00×10^−6^). Theta peak-to-trough was overall lower (prefrontal cortex: 4.95±0.01; reuniens: 5.19±0.01; hippocampus: 11.22±0.02) than beta peak-to-trough (prefrontal cortex: 19.45±0.02; reuniens: 17.84±0.06; hippocampus: 23.86±0.03). Overall, beta was more coupled to delta in all three regions. However, note that in hippocampus theta was more coupled to beta amplitude compared to prefrontal cortex and reuniens (**Figure S7**). This result is consistent with the observed memory-related prefrontal-hippocampal and reuniens-hippocampal coherence curves (**Figure S4**).

Altogether, the results suggest complex dynamics between the beta and theta band occur during sequence memory in which theta shows phasic increases at the onset and offsets of trials, whereas beta is highest during a trial rising in reuniens first and then simultaneously in prefrontal cortex and the hippocampus. All regions showed varying degrees of beta phase-amplitude coupling with delta, and only the hippocampus and prefrontal cortex phase-amplitude coupled with theta. The coupling in theta and delta may be related to the rising and falling of the beta bursts.

### Stimulating reuniens is sufficient to elicit changes in prefrontal-hippocampal LFP activity

While the latency analysis was consistent with the model that reuniens drives prefrontal-hippocampal synchrony, we wanted to test the *causal* capability of reuniens in generating beta prefrontal-hippocampal coherence in freely-behaving rats. Thus, we tested whether reuniens activity was sufficient to drive prefrontal-hippocampal beta synchrony using a retrograde optogenetic viral approach (retroAAV-ChR2-GFP). This approach allowed us to stimulate retrogradely-labeled neurons in reuniens that project to hippocampus (see methods), including a population of dual prefrontal-hippocampal projecting cells (Viena *et al*., 2021). While stimulating, we recorded the LFP activity in both prefrontal cortex and hippocampus (0.002” diameter stainless-steel wires; Allen *et al*., 2020; **Figure 4A_i_**, S1C) as rats freely explored an open-field. There were no memory demands in this experiment.

**Figure 4.**
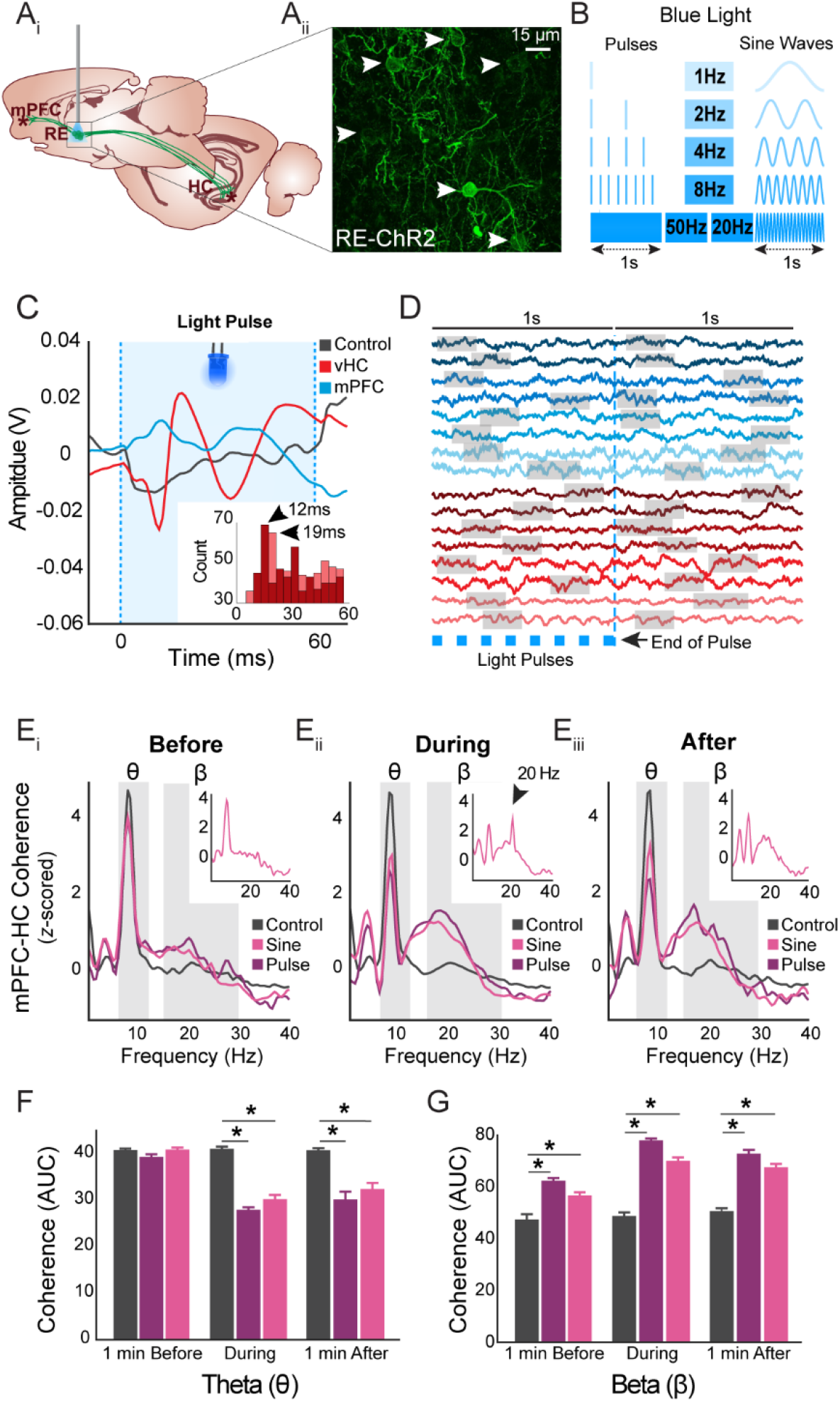
Optogenetic stimulation of reuniens drives beta coherence in prefrontal-hippocampal system. **A**_**i**_) Bilateral injections of retrograde pAAV-Syn-ChR2(H134R)-GFP (experimental) or AAV-CAGGFP (controls) were delivered in vCA1 for the retrograde expression of RE neurons. An optic fiber above reuniens stimulated channelrhodopsin or fluorescence protein expressing neurons. **A**_**ii**_) Confocal image (60x) showing retrograde expression of ChR2 in RE neurons (indicated by white arrows). **B)** LED based blue light pulse or sinewave stimulations at 1,2,4,8, 20 or 50 Hz were used to activate RE-ChR2 transduced and control rats. **C)** Averaged evoked responses in ventral hippocampus resulting from reuniens 8 Hz pulse blue light stimulations (60 ms each) in a control and an experimental subject. As illustrated, stimulating RE-ChR2 expressing neurons resulted in two consecutive negative deflections suggestive of a direct monosynaptic response in vCA1 (in red). This effect was not seen in controls (black). Mild to moderate LFP changes in prefrontal activity were also observed (blue) which may be the result of stimulating putative reuniens mPFC-HC dual projecting neurons expressing ChR2. **D)** Sample averaged raw traces in prefrontal cortex (blue) and hippocampus (red; two samples from each experimental rat) showing a predominant beta rhythm one second prior and one second after the last optogenetic pulse (two samples from each rat). Gray boxes highlight areas of high beta activity. **E)** *z*-scored mPFC-HC coherence in RE-ChR2 and controls across three time points: **E**_**i**_) one min before where all rats have similar theta and beta amplitude values, **E**_**ii**_) during blue light stimulation where there are marked increases in the theta and beta band and a decrease in theta in experimental rats but not controls, and **E**_**iii**_) one min after stimulations where coherence is similar to **E**_**ii**_. Notably, stimulating RE-ChR2 neurons with a 20 Hz sine resulted in a large increase in beta (as previously described) but with an additional large peak around the 20Hz frequency that surpassed the amplitude of delta and theta bands (**E**_**ii**_ inset, black arrow), this effect was not seen before or after stimulations (**E**_**i &**_ **E**_**iii**_ inset). **F)** Theta AUC coherence was significantly different across conditions (control, pulse, sine) and time windows (before, during, after). Notably, theta coherence in control was similar across all time points. **G)** Beta AUC coherence was significantly different across conditions and time windows. In controls, beta coherence was similar across all time points in controls. Abbreviations: AUC, area under the curve; ChR2, channelrhodopsin, HC, hippocampus; min, minute; ms, milliseconds; mPFC, medial prefrontal cortex; RE, reuniens; vCA1, ventral CA1.

Specifically, we used a retrograde adenoassociated virus (retroAAV-ChR2-GFP) to deliver channelrhodopsin to reuniens neurons (ChR2: experimental group) through injections into ventral slm of the hippocampus and compared these to matched retroAAV-GFP controls. We confirmed expression patterns histologically (post-mortem) with co-expressed GFP markers using fluorescence confocal microscopy (**Figure 4A_ii_**). We found ChR2 expressing cells in reuniens near the optrode tips in all ChR2 rats, including GFP-labeled axons leaving reuniens, prefrontal cortex and hippocampal slm sites, confirming the reuniens ChR2 expression targets and circuits. Blue light was delivered via an optic fiber positioned above reuniens with an LED source in pulse (1, 2, 4, 8 and 50 Hz) or sine wave patterns (1, 2, 4, 8, 20 Hz; **Figure 4B**). The use of multiple frequencies helped determine if a specific stimulus pattern was needed to induce prefrontal-hippocampal synchrony.

We first assessed whether the blue light had any effect on the LFP activity in prefrontal-hippocampal recording sites across days and groups (**Figure S8**). We quantified and averaged 1 s periods of raw voltage signals relative to the onset of the 8 Hz blue light pulses and to 4, 8 and 20 Hz sinewaves (**Figure S8**) in order to confirm this approach evoked responses in downstream regions, as anticipated, and also to compare the efficacy of our stimulation patterns. In ChR2 rats, we observed that blue light delivered to reuniens evoked strong monosynaptic responses in ventral hippocampus (**Figure 4C**). This response was in line with other reports of reuniens-evoked ventral hippocampus responses (Dolleman-van der Weel, Lopes da Silva and Witter, 1997); **Figure 4C, S8A-B**). We replicated this effect on multiple days in the ChR2 rats, but importantly, we never observed evoked potential in controls. This demonstrated the specificity of the effect of ChR2 activation in reuniens (**Figure 4C, S8A-B**). We also observed similar latency, but much less pronounced, evoked activity in prefrontal cortex. This was consistent with the retroAAV approach leading to ChR2 expression in a smaller populations of dual hippocampal-prefrontal projecting reuniens neurons (Hoover and Vertes, 2012; Viena *et al*., 2021).

We then measured the latency of each raw voltage trace negative and positive peak occurrences in ventral hippocampus following the onset of each pulse stimulation. As shown in inset **Figure 4C** (negative peaks in red, positive peaks in pink) during the 60 ms pulse duration, the frequency of the negative peaks was highest during the 10-15 ms bin (first negative peak) and then at 30-35 ms (second negative peak), while the large positive peaks appeared between 15-20 ms (first positive peak) and 50-55 ms (second peak). The modal latency of negative peaks during pulses was 12 ms, indicating that our reuniens optogenetic manipulations elicited direct monosynaptic response in ventral hippocampus, consistent with the anatomy and physiology of reuniens glutamatergic projections into the hippocampus (Hoover and Vertes, 2007; Vertes *et al*., 2007; Viena *et al*., 2021).

The sine wave stimulations also caused changes to the voltage activity recorded in prefrontal and ventral hippocampus (in ChR2 rats only). However, the max-to-min voltage amplitudes were 2-10x smaller than the pulse evoked responses (**Figure S8C**). Sine wave stimulations triggered rapid down-state transitions at the rise of the sinewave, and rapid up-state transitions with the fall of the sine wave. That is, sinusoidal stimulations essentially created weak rhythmic voltage activity in both prefrontal and hippocampal traces, albeit a bit more square-shaped than the input stimulus.

### Reuniens drives prefrontal-hippocampal beta coherence

We next looked if reuniens induced prefrontal-hippocampal coherence at larger time scales (seconds-to-minutes). The raw traces showed the presence of delta, theta and beta rhythms during and after the stimulation and generally looked similar to recordings from the memory experiments (see example traces in **Figure 4D**, highlighted with gray boxes). We observed delta and theta rhythms in prefrontal cortex and hippocampus in ChR2 and control rats, with more delta in prefrontal sites and more theta in hippocampal sites. However, beta was only commonly seen in traces from ChR2 rats after stimulation with blue light (experienced recently or in another session). This suggested reuniens activation may be sufficient to increase beta in prefrontal cortex and hippocampus.

Next, we quantified and tested the capacity of reuniens neurons to drive beta and theta coherence in the prefrontal-hippocampal network. We assessed coherence one minute before, during and after the end of stimulations. Surprisingly, we found that prefrontal-hippocampal coherence curves were largely independent of the stimulation pattern. See **Figure S9** for all the coherence curves calculated in ChR2 rats by stimulus type (pulsed vs. sine), time (1 min before, during and after), frequency (1, 2, 4, 8, 20, and 50 Hz) and under the same conditions in controls.

For quantification, we collapsed all frequencies of the pulse and sine stimulations together into mean group coherences shown in **Figure 4E**. One minute before the onset of stimulations, coherent activity between prefrontal and hippocampus was similar in ChR2 and control rats (**Figure 4E_i_**). During reuniens stimulations, coherence between prefrontal cortex and hippocampus changed in ChR2 rats compared to controls in a band specific manner. Theta coherence showed a significant reduction, and beta coherence showed a significant increase. The beta coherence increase was a large effect (∼2 at the peak) and broad from ∼10-30 Hz (**Figure 4E_ii_**). Interestingly, beta coherence remained high for at least one minute after stimulations had terminated in both pulse and sine, however, we did not see this residual beta effect in control rats which also received blue light pulse stimulations (**Figure 4E_iii_**), thus eliminating the observed beta as a simple artifact of the blue light stimulations. Notably, this residual beta coherence was somewhat evident in both pulse and sine in the one minute before coherence plot (**Figure 4E_i_**). It is important to point out that the majority of stimulation blocks included in the analysis had been preceded by other reuniens blue-light stimulated blocks (see Methods). Notwithstanding, the change in coherence from one minute before, to during, and one minute after was significantly different in experimental subjects compared to controls (see stats below). One noticeable effect was a 20 Hz sine wave caused a narrow 20 Hz prefrontal-hippocampal coherence peak (inset in **Figure 4E_ii_**), riding on top of the broader beta band coherence induction, suggesting that the network was particularly resonant within the beta band frequencies. Optogenetic stimulation of reuniens increased beta coherence, and decreased theta coherence in a pattern closely matching what was observed in the memory experiments (**Figure 1F_i_**). In control rats, prefrontal-hippocampal coherence was always dominated by theta with little-to-no beta observed.

For inferential comparisons, we calculated the AUC for our *a priori* theta and beta bands across the three different time windows (one minute before, during, one minute after) and conditions (control, pulse, sine) **(Figure 4F)**. There was a significant effect of time across all time points on theta coherence (F_(2,36)_ = 58.08, *p* = 5.39×10^−12^) with significant reductions between theta_before_ and theta_during_ (*p* = 1.30X10^−11^), theta_before_ and theta_after_ (*p* = 2.50×10^−9^), but no difference between theta_during_ and theta_after_ (*p* = 0.19). We also found a main effect of condition (F_(2,36)_ = 75.07, *p* = 1.43×10^−13^) with significant reductions in theta_control_ and theta_pulse_ (*p* = 2.14×10^−13^), in theta_control_ and theta_sine_ (*p* = 3.87×10^−10^), but not difference between theta_pulse_ and theta_sine_ (*p* = 0.72). We also found a significant time x condition interaction effect (F_(4,36)_ = 15.18, *p* = 2.32×10^−7^) indicating that RE activation overall decreases prefrontal-hippocampal coherence in the theta band in ChR2 rats during and after stimulations compared to controls. We also found a significant effect of time across all time points on beta coherence (**Figure 4G**; F_(2,36)_ = 27.24, *p* = 6.25×10^−8^) with significant increases in beta coherence between beta_before_ and beta_during_ (*p* = 1.77×10^−7^), beta_before_ and beta_after_ (*p* = 3.00 ×10^−6^), but no difference between beta_during_ and beta_after_ (*p* = 1.00). We also found a main effect of condition (F_(2,36)_ = 149.82, *p* = 3.53×10^−18^) with significant increases in beta_control_ and beta_pulse_ (*p* = 2.19×10^−18^), in beta_control_ and beta_sine_ (*p* = 1.86×10^−12^), and between beta_pulse_ and beta_sine_ (*p* = 1.00×10^−6^). We also found a significant time x condition interaction effect (F_(4,36)_ = 8.15, *p* = 8.7×10^−5^) indicating that reuniens activation overall increased beta coherence in ChR2 rats during and after stimulations compared to controls. This shows that optogenetic activation of reuniens significantly increases prefrontal-hippocampal beta coherence while it decreases theta coherence in ChR2 rats during and after stimulations, but not in controls.

Altogether, these results demonstrate that reuniens activation is sufficient to drive prefrontal-hippocampal beta coherence modes but reduce theta coherence. Importantly, this pattern of decreased theta and increased beta coherence during reuniens optogenetic stimulations was similar to the pattern observed in prefrontal-hippocampal coherence during the nonspatial sequence memory task.

## DISCUSSION

### Summary of main findings

In the first experiment we recorded LFPs using dual-site silicon probes targeting the medial prefrontal cortex and hippocampus during performance on a nonspatial sequence memory task. We found that memory was related to a transient ∼15-30 Hz beta mode, which accounted for a plurality of the coherence between the prefrontal cortex and hippocampus sites during memory trials. In the second experiment, we recorded LFPs using dual-site silicon probes targeting the nucleus reuniens of the thalamus and hippocampus during performance on the same nonspatial sequence memory task. Results revealed that high amplitude beta bursts appear in reuniens shortly after a trial begins, and shortly before beta onsets in the hippocampus. This suggests that reuniens might help drive the synchronous beta observed in the prefrontal-hippocampal recordings. We then combined data from both experiments to scrutinize beta time courses across the entire prefrontal-reuniens-hippocampal memory system. This analysis provided evidence that beta first appears in reuniens and is then followed by the simultaneous and concurrent rising of beta in the prefrontal cortex and hippocampus, highly suggestive of a reuniens to prefrontal-hippocampal directionality. In our third experiment, we used an optogenetic approach (retroAAV-ChR2) to test if reuniens cells can drive this type of prefrontal-hippocampal synchrony in naïve rats navigating an open field. Stimulating reuniens reliably increased prefrontal-hippocampal beta coherence, and decreased theta coherence, in a pattern resembling the memory mode observed during the sequence task. The results from all three experiments provide strong evidence that reuniens activity drives prefrontal-hippocampal synchrony and support a role for reuniens in orchestrating a prefrontal-hippocampal beta operation mode conducive to using long-term memory stores to help guide current situational demands.

### Overt behavior determines rapid transitions in rhythmic prefrontal-hippocampal interactions modes

Not surprisingly, the prefrontal-hippocampal interaction modes observed were dynamic and dependent on the overt behavioral state of the animal (Buzsáki and Eidelberg, 1983; Hyman *et al*., 2005). Our recordings show that novel beta-dominated interaction modes came on rapidly (< 1 s) during trials of a nonspatial sequence memory task, and are interleaved between theta-dominated interactions modes driven by running behaviors (Maurer *et al*., 2005; Hinman *et al*., 2016). Prefrontal-hippocampal coupling was generally weaker across frequencies when the rat was not engaged in either behavior, likely meaning this system is less integrated at these times, or in modes defined by behaviors not well separated here (e.g., awake delta modes, Schultheiss *et al*., 2020).

### Prefrontal-hippocampal beta modes reflect memory

We performed analyses suggesting prefrontal-hippocampal beta coherence reflects the cognitive aspects of the sequence memory task, and not simply sensory aspects of the odor processing or the poking behaviors alone. This analysis showed accurate in sequence trials had significantly stronger beta coherence than inaccurate out of sequence trials. Thus, prefrontal-hippocampal beta reflected the mnemonic aspects of the sequence memory task, and not simply the odors or poking behaviors. This conclusion was bolstered by an additional analysis of the local beta power in the prefrontal cortex and hippocampus regions separately. This showed significant activity in both regions during accurate in sequence trials, but never showed significant activity on inaccurate out of sequence trials. This suggests that beta activity in each region separately also reflect mnemonic aspects of the task, and consistent with the notion that their shared activity would also reflect mnemonic aspects of the task.

These results add to the field by extending the scope of the known cognitive beta networks of rodents to include prefrontal-hippocampal interactions in memory, and specifically nonspatial memory for sequences of events (Jayachandran *et al*., 2019). Along these lines, a previous study found beta coherence between the olfactory bulb and hippocampus was increased when transferring a go/no-go rule to a new odor set, but not present for simple odor learning concluding that beta coherence was related to the flexible use of context appropriate odor memories (Martin, Beshel and Kay, 2007). Notably, rodent prefrontal cortex is critical to learning multiple lists of odors and resolving interference (Peters *et al*., 2013). In the sequence task multiple odor lists are repeatedly and concurrently presented InSeq or OutSeq, in which knowledge of the temporal contexts alone can resolve the appropriate response. Other studies have shown beta coherence between the lateral entorhinal cortex and hippocampus occurring during memory for odor place associations, which are thought to reflect the learning of navigational odor associations (Igarashi *et al*., 2014). Lastly, our data are also consistent with the description of memory-related hippocampal beta power found in single list odor sequence memory (identified as 20-40 Hz) showing hippocampal neurons entrain to beta, code for sequential context of odors (Allen *et al*., 2016), and may be strongest in proximal CA1 (Gattas *et al*., 2022). The present data from a two-list odor sequence task the beta activity in those studies probably also reflects prefrontal-hippocampal interactions, and thus was part a larger integrated memory and decision-making network.

### Reuniens-hippocampal interaction modes transition rapidly dependent on behavior

Our recordings also found that reuniens, a midline thalamic structure that bidirectionally connects the prefrontal cortex and hippocampus (Vertes *et al*., 2006; Vertes, Linley and Hoover, 2015; Dolleman-van der Weel *et al*., 2019), shows similarly high coherence with the hippocampus. Like with prefrontal cortex, these modes dynamically shift with behavior. In the reuniens-hippocampal coherence analysis we also observed beta-dominated interaction modes during a nonspatial sequence memory task, interleaved between theta-dominated interactions modes driven by running (Maurer *et al*., 2005; Hinman *et al*., 2016). This was hypothesized, but not known, based on the anatomical work showing that glutamatergic reuniens neurons form a large portion of the synapses in slm of the CA1 including contacts on both pyramidal cells and GABAergic interneurons (Dolleman-van der Weel, Lopes da Silva and Witter, 1997; Dolleman-van der Weel and Witter, 2000). In this latter regard, reuniens is thought to provide powerful feed-forward inhibition resulting in changes in the local excitatory-inhibitory balance (Dolleman-van der Weel *et al*., 2019), and thus capable of setting modes or pacing the system. Notably, reuniens has previously been shown critical to performance-related theta-gamma prefrontal-hippocampal synchrony in a spatial working memory task (Hallock, Wang and Griffin, 2016). In line with this coupling, we observed a theta-dominated reuniens-hippocampal coherence mode during running.

However, reuniens-hippocampal beta modes have not been previously reported. Reuniens-hippocampal beta modes were in line with our prefrontal-hippocampal beta modes in terms of having onsets a half sec into trials, high coherence values, and a defining ∼15-30 Hz frequency range. Unique to reuniens, we found a different pattern of significance when looking across the sequential trial types of the sequence task. That is, we found no significant differences between trials with similar poke times. While not definitive in ruling out a mnemonic role for reuniens-hippocampal beta, or reuniens itself, this pattern might instead reflect a general role for reuniens in engaging prefrontal-hippocampal interactions. In line with this analysis, only reuniens recordings showed significant beta power on out of sequence incorrect trials, and thus did not strongly differentiate different sequence memory trials with similar poke times. It is worth pointing out here that while reuniens efferents are primarily restricted to medial prefrontal cortex, CA1 and lateral entorhinal cortex, its afferents come from a very a large number of regions including the brainstem, amygdala, hypothalamus, thalamic reticular nucleus, and limbic cortex and more. This places reuniens in position to integrate internal states with other behavior-related activity into a “go memory network” signal at the appropriate times held on through the duration of the need for memory (Dolleman-van der Weel *et al*., 2019). Viewed in this way the reuniens may be driven by the behavioral situation (the sequence task), and then orchestrate the activation of the prefrontal-hippocampal memory network in response to aid in decision-making.

### The times course of beta in the prefrontal-reuniens-hippocampal system

The time course of beta found here is consistent with a role in engaging the memory network. Several temporal results are relevant here including: (1) Reuniens beta had a significant probability of coming on ∼100 ms before the animal started a trial, thus precluding that it is triggered by the odor itself on those trials and suggests the response is instead anticipatory. (2) Reuniens beta preceded prefrontal-hippocampal beta but with short latencies. This is consistent with the dual-site monosynaptic projections from reuniens neurons. (3) Prefrontal-hippocampal beta came on simultaneously, as predicted by reuniens efferents. (4) Prefrontal-reuniens-hippocampal beta rose together to maximal levels before a decision was made, after such time that sequence cells in CA1 are known to activate (Allen *et al*., 2016). This suggests they may help maintain informational stasis (e.g., Lundqvist *et al*., 2016). (5) Beta dropped briefly just before the decision. This suggests beta releases in time to release successful behaviors. (6) Prefrontal-reuniens-hippocampal beta collapsed to baseline levels shortly after the decision and in unison. This time course would also be consistent the finding that reuniens plays a role in working memory because it vanishes after a temporary use period (Hallock, Wang and Griffin, 2016; Viena, Linley and Vertes, 2018; Jayachandran *et al*., 2019). Additionally, the coherence was transitioned rapidly enough and was brief enough to be involved in repeated top-down directed memory retrievals, as is necessary in the sequence memory task. This model of the cognitive timeline for beta in working memory will require future causal tests that can interrupt beta at very precise times, and an examination of the mnemonic content, such as may be afforded by optogenetics or electrical stimulation and ensemble analysis.

### Beta comes in bursts and couples to local delta and theta phases

Peak prefrontal-hippocampal beta coherence magnitudes were very high during memory trials, at least equal in magnitude to those obtained in theta during running. The suggests activity in these areas were precisely timed for long-range integrations (Fries, 2005). The mechanisms of beta coherence are unknown, but undoubtedly involve well-timed local inhibitory activity and cross-frequency coupling with lower frequencies (Buzsáki *et al*., 2003; Canolty *et al*., 2006). That is, beta may reflect the rhythms of the local region’s excitatory-inhibitory tone, but the timing of bursts may be shaped by the phases of delta or theta, similar to theta-gamma couplings (Tort *et al*., 2009; Belluscio *et al*., 2012). Here, we show that beta amplitudes were strongly coupled to delta (1-4 Hz) waves throughout prefrontal-reuniens-hippocampal system. Additionally, inspection of the prefrontal-hippocampal coherence curves show that high magnitude delta coherence often coincides with beta coherence during the sequence task. Thus, from an interregional synchrony standpoint, ‘beta modes’ might more accurately be called ‘delta-beta modes’. Notably, beta amplitudes also coupled to theta phases, but only in hippocampal sites (consistent with Igarashi *et al*., 2014). Along this line, beta was busty and transient, coming in amplitude waves that could be described as resembling “beat frequencies” lasting ∼100-240 ms. This pattern seemed consistent with potential amplitude modulations in the delta range (Schultheiss *et al*., 2020). Along these lines, reuniens has a known role in coordinating prefrontal-hippocampal networks during slow wave sleep and in a state dependent manner (Hauer, Pagliardini and Dickson, 2019, 2022).

### Reuniens activity alone is sufficient to drive prefrontal-hippocampal beta synchrony and reduce theta synchrony

Can reuniens synchronize prefrontal-hippocampal networks? There is a growing body of literature from anesthetized sleep-like states that reuniens can help synchronize networks, especially at delta or slower frequencies (Ferraris *et al*., 2018; Hauer, Pagliardini and Dickson, 2019). However, little is known about the awake behaving reuniens. Using an optogenetic approach, we showed that stimulating reuniens lead to large monosynaptic responses in the hippocampus that resulted in prefrontal-hippocampal increased beta and decreased theta coherence that resembled the novel beta coherence state that was driven by the sequence memory task. The resulting coherence is likely due to increased feed-inhibition in CA1 (Dolleman-van der Weel, Lopes da Silva and Witter, 1997).

It is well known that cortico-thalamo-cortical circuits are essential in coordinating activity across cortical brain regions. Beta synchronization between cortical regions with or via the thalamus is plausible. For instance, mediodorsal nucleus has been shown to synchronize with prefrontal cortex in the beta range (13-30 Hz) during acquisition and performance of a working memory task (Parnaudeau *et al*., 2013) and unilateral ablation of ventral medial thalamus appears to be responsible for the exaggerated high beta range oscillatory activity between the basal ganglia-thalamo-motor cortex network (Brazhnik *et al*., 2016). Our findings are in line with thalamic nuclei being able to influence cortical responses by increasing beta activity in distal regions, and/or adjust the degree of synchrony of these regions to enhance thalamo-cortical transmission as supported by our data.

### Do beta bursts contribute to episodic buffers in working memory?

Although beta has been widely studied for movement and odor processing, the time course of beta observed here is similar to cortical beta burst profiles reported in non-human primates in working memory (Lundqvist, Compte and Lansner, 2010; Lundqvist *et al*., 2016). More specifically, modulations of beta rhythmic activity in non somato-motor areas have been associated with decision-making (Pesaran, Nelson and Andersen, 2008; Wimmer *et al*., 2016). Along this line, studies have consistently found that beta displays cognitive roles during odor tasks (Martin *et al*., 2006; Igarashi *et al*., 2014; Allen *et al*., 2016; Gattas *et al*., 2022). Here, we show that beta is transiently engaged when episodic-like memory is needed to make a successful decision. Such content-specific beta activity has been observed during endogenous information processing in working memory and decision-making (Buschman *et al*., 2012; Salazar *et al*., 2012; Lewis *et al*., 2016; Wimmer *et al*., 2016). We also show that beta activity reflects the sequential content of a trial and is not strictly driven by sensory or motor cues since prefrontal-hippocampal coherence occurs after the odor is presented. Finally, we provide evidence that beta can be generated internally in reuniens with optogenetic activations. In other words, beta generated from reuniens may then influence the prefrontal-hippocampal system to hold relevant information (temporary storage) within a short period of time (beta burst) to make a decision. Together, this leads one to speculate if reuniens-related beta modes could be part of the neurobiological basis of an episodic buffer in working memory (see Baddeley, 2000).

## Conclusion

The present study tested the hypothesis that network-wide prefrontal-reuniens-hippocampal synchrony is engaged in a nonspatial sequence memory task reflecting an intrinsic connectivity network state conducive to memory-based decision making. We also tested the prominent hypothesis that the nucleus reuniens of the thalamus contributes to memory by providing synchronizing drive to the medial prefrontal cortex and the hippocampus (Eichenbaum, 2017; Dolleman-van der Weel *et al*., 2019). We provide several new insights into the neurobiological underpinnings of episodic memory, and direct evidence for the prevalent theory that thalamic reuniens synchronizes prefrontal and hippocampal networks at appropriate times, allowing the flexible use of memory to benefit situation-specific behaviors (e.g., using episodic memories to help guide decisions; Eichenbaum, 2017).

## Acknowledgements

This work was supported by National Institutes of Health (NIH) grants R01 MH113626 and F99 NS119001 and FIU. A special thanks to all the members of the Allen Lab, specifically our undergraduate research assistants, F.P.R., N.T., G.E.R., D.S., N.L., C.R., K.K.R., and J.M., who helped with task training and data collection, pre-processing and histology. We would like to thank the Animal Care Facility and Dr. Horatiu Vinerean.

## Author Contributions

Conceptualization, M.J., T.D.V., and T.A.A.; Methodology, M.J., T.D.V., and T.A.A.; Investigation, M.J. and T.D.V., A.G., A.V.V, S.L, and V.R; Writing – Original Draft, M.J., T.D.V., and T.A.A.; Writing – Review and Editing, T.A.A., M.J., T.D.V., A.G., A.V.V., S.L., V.R., and R.P.V.; Funding Acquisition, T.A.A., R.P.V., T.D.V., and M.J.; Resources, T.A.A., R.P.V.; Supervision, T.A.A.

## METHODS

### Subjects

19 Long-Evans rats (10 females) from Charles River Laboratories weighing 250-350 g upon arrival were used. Rats were individually housed and maintained on a 12-h inverse light/dark cycle (lights were turned off at 10 AM). Rats trained on the sequence memory task had *ad libitum* access to food and enrichment, but access to water was limited to 3 – 5 min each day, depending on how much water they received as a reward during behavioral training (6 – 9 mL). Rats used in the open field task had ad libitum access to food, enrichment, and water. All training and testing sessions were conducted during the dark phase (active period) of the light cycle. All experimental procedures using animals were conducted in accordance with the Florida International University Institutional Animal Care and Use Committee (FIU IACUC). Although we used male and female rats, we did not evaluate sex as a variable since we have previously demonstrated there are no differences in performance between males and females on the sequence memory task (Jayachandran *et al*., 2022).

### Sequence Memory Experiments

#### Sequence Memory Task

The sequence memory task used (Allen *et al*., 2014, 2016; Jayachandran *et al*., 2019, 2022) involves repeated presentations of odor sequences and requires the rat to determine whether each item (odor) was presented in sequence (InSeq; by holding the nose-poke response for 1 s) or out of sequence (OutSeq; by withdrawing its nose from the port before 1 s). Rats were trained on two sequences, each comprising four distinct odors (e.g., Seq1: ABCD, Seq2: WXYZ). Each sequence was presented at either end of a linear track maze. Odor presentations were initiated by a nose-poke, and each trial was terminated after the rat either held the nose-poke response for > 1 s (InSeq) or withdrew its nose-poke response for < 1 s (OutSeq). There was a 1 s interval between trials. Water rewards (20 μl; diluted at 1 g of aspartame for every 500 mL of water) were delivered below the odor port after each correct response. Following an incorrect response, a buzzer sound was emitted and the sequence was terminated. Each sequence was presented alternatively 50-100 times per session; approximately half the presentations included all items InSeq (ABCD) and half included one item OutSeq (e.g., AB<u>A</u>D, odor A repeated in the 3rd position). Note OutSeq items could be presented in any sequence position except the first position (i.e., sequences always began with an InSeq item). Sequence memory was probed with OutSeq trials (e.g., AB<u>A</u>D; one OutSeq trial randomly presented per sequence).

#### Sequence Task Apparatus

Rats trained on the sequence memory task were tested in a noise-attenuated experimental room. The behavioral apparatus was comprised of a linear track (length, 183 cm; width, 10 cm; height, 43 cm) with walls angled outward at 15° and nose ports at each end through which repeated deliveries of multiple distinct odors could be presented. Photobeam sensors were used to detect nose port entries. Each nose port was connected to an odor delivery system (Med Associates). Odor deliveries were initiated by a nose-poke entry and terminated either when the rat withdrew before 1 s, or after 1 s had elapsed. Water ports were positioned under each nose port for reward delivery. Timing boards (Plexon) and digital input/output devices (National Instruments) were used to measure all event times and control the hardware. All aspects of the task were automated using custom MATLAB scripts (MathWorks R2016a). A 256-channel Omniplex D with video tracking and Cineplex behavior software (Plexon) were used to interface with the hardware in real time and record behavioral data. Odors were organic odorants contained in glass jars (A: 1-octanol; B: (-) - limonene; C: I-menthone; D: isobutyl alcohol; W: acetophenone; X: (1S) - (-) – beta pinene; Y: L (-) - carvone; X: 5-methyl-2- hexanone) that were volatilized with nitrogen air (flow rate, 2 L/min) and diluted with ultrapure air (flow rate, 1 L/min). To prevent cross-contamination, separate Teflon tubing lines were used for each odor. These lines converged into a single channel at the bottom of the odor port. In addition, a vacuum located at the top of the odor port provided constant negative pressure to quickly evacuate odor traces with a matched flow rate.

#### Sequence Memory Task Training

Naive rats were initially trained in a series of incremental stages over 20-30 weeks. Each rat was trained to poke and hold its nose in an odor port to receive a water reward. The minimum required nose-poke duration started at 50 ms and was gradually increased (in 15 ms increments) until the rat reliably held the nose-poke position for 1.2 s for ∼ 70% of the time over three sessions (75-100 nose-pokes per session). The rats were then habituated to odor presentations in the port (odor A_1_ and A_2_, then odor sequences A_1_B_1_ and A_2_B_2_) and each rat was required to maintain its nose-poke response for 1 s to receive a reward. The rats were then trained to identify InSeq and OutSeq items. Rats were initially trained on a two-item sequence: they were presented with ‘‘AB’’ and ‘‘A<u>A</u>’’ sequences in equal proportions. The correct response to the first odor was to hold the nose-poke for 1 s (Odor A was always the first item). For the second odor, rats were required to determine whether the item was InSeq (AB; hold for 1 s to receive reward) or OutSeq (A<u>A</u>; withdraw before 1 s to receive a reward). After reaching criterion on the two-item sequence, the number of items per sequence was increased to three and four in successive stages (criterion: ∼60% correct across all individual odor presentations over three sessions). After reaching criterion performance on the two four-odor sequences (60% correct on both InSeq and OutSeq items), rats underwent surgery to implant chronic recording electrodes.

#### Prefrontal-Hippocampal Implants

Five rats were implanted with 32-channel silicon probes arranged as tetrodes in prefrontal cortex and hippocampus with 25 μm between adjacent electrode sites and impedances of 1.23 ± 0.32 MΩ. The eight tetrodes were distributed across four shanks at tip-to-tetrode depths of 78 μm and 228 μm (NeuroNexus A4X2-tet-5mm). Shanks were separated by 200 μm giving each probe a total length of 0.67 cm. During implantation, the long axis was oriented medial-lateral. One rat was implanted with one silicon probe targeting prefrontal cortex (A/P 3.24 mm, M/L 0.6-1.2 mm, D/V_from cortex_ -3.45 mm). Four rats were implanted with silicon probes targeting both prefrontal cortex (A/P 3.24 mm, M/L 0.6-1.2 mm, D/V_from cortex_ -3.45mm) and hippocapus (specifically CA1; A/P -3.24 mm, M/L 2.4-3.0 mm, D/V_from cortex_ -2.4 mm). An additional four rats were implanted with 32-channel silicon probes in reuniens and hippocampus with impedances of 0.043 MΩ. The 32 single electrodes were distributed across two shanks spanning a length of 300μm from the tip (Cambridge NeuroTech AASY-116 E-1 & E-2). Shanks were separated by 250 μm. During implantation, the long axis was oriented medial-lateral. Four rats were implanted with silicon probes targeting both reuniens (A/P 1.92 mm, M/L 0.3-0.55 mm, D/V_from cortex_ 7.0 mm) and hippocampus (specifically CA1; A/P 3.24 mm, M/L 2.4-2.65 mm, D/V_from cortex_ 2.5 mm)

#### Prefrontal-Hippocampal and Reuniens-Hippocampal Surgery

Rats were anesthetized with isoflurane (induction 5%; maintenance: 2-3%) mixed with oxygen (800 ml/min) and placed in a stereotaxic apparatus (David Kopf Instruments, Model 900). A protective ophthalmic ointment (Gentak, 0.3%) was applied to the eyes and the scalp was sterilized with applications of isopropyl alcohol (70% in deionized H_2_O) followed by Betadine. The incision site was locally anesthetized with Marcaine® (7.5 mg/ml, 0.5 ml, s.c.) and the skull was exposed following a fisheye incision. Adjustments were made to ensure that bregma and lambda were level (±0.05 μm in the D/V plane). Body temperature (35.9-37.5ºC) was monitored and maintained throughout surgery using a rectal thermometer and circulating water heating pad. Ringer’s solution with 5% dextrose was administered to maintain hydration (5 ml, s.c.), and glycopyrrolate (0.2 mg/ml, 0.5 mg/kg, s.c.) was administered to prevent respiratory difficulties.

For silicon probe implants using the RatHat (Allen *et al*., 2020) targeting prefrontal cortex (n = 1), a rectangular craniotomy was drilled to accommodate the probe shanks centered on coordinates AP 3.24 mm, ML 0.9 mm. For dual-site implants in prefrontal cortex and hippocampus (n = 4), two craniotomies were drilled targeting prefrontal cortex (AP 3.24 mm, ML 0.9 mm) and hippocampus (at AP −3.24 mm, ML 2.7 mm). For dual-site implants using the RatHat (Allen et al., 2020) targeting reuniens and hippocampus (n = 4), two craniotomies were drilled targeting reuniens (A/P 1.92 mm, M/L 0.3-0.55 mm, D/V_from cortex_ 7.0 mm) and hippocampus (specifically CA1; A/P 3.24 mm, M/L 2.4-2.65 mm, D/V_from cortex_ 2.5 mm). Burr holes were drilled and skull screws (1/8-inch grade 2 (CP) titanium; Allied Titanium Inc) were secured onto the skull. After removal of the dura in the craniotomy, implants were lowered on the stereotaxic arm until the electrode tips were just above the cortical surface. The ground wire was attached to the ground screws, and implants were lowered such that prefrontal cortex electrodes reached ∼3.3 mm below the cortical surface and hippocampus electrodes reached ∼2.4 mm in depth. 0.5% sodium alginate solution was applied with a syringe and 23 ga. needle onto the exposed brain tissue within the craniotomy, then several drops of a 10% calcium chloride solution was applied to fix the alginate into a gel. The silicon probe was affixed to the surgical screws with dental cement (methyl, methacrylate). A protective cover was created using a high-resolution (56 mm) stereolithography 3D printer (ProJet 1200; 3D Systems), suitable for chronic head stages. A custom-designed 3D-printed protective wall and cap was created using CAD software (Autodesk Inventor Pro Edition) and assembled around the exposed silicon probe to protect the headstage against impact and debris. Excess skin was sutured (black silk suture 4-0, with reverse cutting needle 19 mm, 1/2 Circle). Neosporin® was applied to the skin surrounding the head stage. At the end of surgery, Flunixin (50 mg/ml, 2.5 mg/kg, s.c.), a nonsteroidal anti-inflammatory analgesic, was administered to the rats. The rats were returned to a clean cage and monitored until they awoke. One day following surgery, the headstage was checked, the rats were administered a dose of Flunixin, and Neosporin® was reapplied. At least one week was allowed for recovery from surgery prior to beginning experiments. Rats with incorrect placement of silicon probes or defective electrodes (*n* = 0) were not considered in the analysis.

#### Prefrontal-Hippocampal and Reuniens-Hippocampal Electrophysiological Recordings

Rats were recorded during the sequence memory task for three consecutive sessions. Throughout each experiment wide-band data was acquired with digital headstages (32-channel, 40kHz sampling rate) and OmniplexD systems (256-channel, Plexon) coupled to an automated behavioral rig. The OmniplexD systems acquires 32 digital behavioral event inputs, 16 analog event inputs, and digital video (80fps). Voltage signals recorded from the tetrode tips were referenced to a ground screw positioned over the cerebellum (low cutoff = 0.7 Hz). Local field potentials (LFP) were separated into a second data stream (LFP: 0.7 Hz–300Hz). LFP frequency bands of interest, including delta (1-4 Hz), theta (6-12 Hz) and beta (15-30 Hz), were filtered offline. LFP activity was analyzed with NeuroExplorer (Plexon) and MATLAB.

### Optogenetic Experiment

#### Reuniens Surgery

Rats were acclimated to the housing facility and handled for at least one week prior to initiation of any surgical procedures. In preparation for surgery, we anesthetized the rats using isoflurane (4-5% induction, 1.5-2% maintenance) and their heads fixed in a stereotaxic apparatus (Kopf). An incision was made on the skin to expose the skull and burr holes were drilled above the prelimbic/infralimbic cortext (A/P: +3.0, M/L: ±0.8); midline thalamus (A/P: -2.0, M/L: -0.2), and ventral hippocampus (A/P: -5.6, M/L: ±5.85) using a RatHat surgical stencil (Allen et al., 2020).

We injected bilaterally 600μL channelrhodopsin-infused retrograde adeno associated virus pAAV-Syn-ChR2(H134R)-GFP (Addgene) in ventral hippocampus (A/P: -5.6, M/L: ±5.85; D/V: -7.4, 200μL, -6.8, 200 μL & -6.2, 100 μL) at a speed of 1.0 nL/sec (Nanoject III, Drummund Scientific).

After injections, the needle was kept in place for an additional 5 minutes to facilitate diffusion. Then, negative pressure was used to avoid upward suction of the virus during the tip retractions from the brain. Upon delivering the viral vector(s), we implanted a custom-built RatHat microelectrode-optrode assembly, described in detail by Allen *et al*., 2020. The RatHat contained stainless steel (SS) wire electrodes that recorded LFP (wire length from skull: prelimbic/infralimbic 5.5-6 mm; and ventral hippocampus 5.5-6 mm). At the conclusion of the surgery, we secured the implant to the skull using titanium bone screws and dental cement. Animals recovered for 3-4 weeks to allow for viral vector expression prior to starting electrophysiological recordings. Microelectrode impedances were tested at four intervals (prior to implantation, one week after surgery, day before testing began, prior to animal’s perfusion) to check for out of range impedances and dead channels. Rats with incorrect placement of optic fiber or SS wires, improper injection site or viral expression, or defective electrodes (*n* = 1) were not considered in the analysis.

#### Reuniens Optic Fiber Placement

Five days prior to commencement of recording sessions, we lightly anesthetized rats for chronic placement of an in-house fabricated optic fiber stub (tip ø: 200 μm, 0.5 NA; Thorlabs). The length of the fiber stub was customized to deliver light intracranially above reuniens (A/P:2.0, M/L: +0.2; D/V length 7.0 mm) via a canula tube in the RatHat implant. We then secured an optic fiber with dental cement. Finally, we tested the light delivery of the optic patch cables along with the optrode and documented/calculated the yield prior to implantation. We ensured the light delivery was within 10–12 mW based on the % power yield using a power meter (Thorlabs) both prior and after each session to make sure light delivery remained stable throughout the experiment.

#### Open Field Behavioral Apparatus

Open field (OF) recordings occurred in a rectangular maze (OF; dimensions: 122 cm x 118 cm x 47 cm) placed 72 cm above the floor (as described in Schultheiss *et al*., 2020). The OF was surrounded by black curtains and room lights were kept off except for red LED lights above the arena.

#### Optical Stimulation

ChR2 expressing cells were optically stimulated using a 465 nm (blue light) compact LED attached to a rotary commutator above the OF (Plexon). The LED was connected to the optic fiber stub implanted in the brain via a patch cable (1.75 m long). Photostimulation events were driven with a PlexBright 4 channel controller and Radiant V2.3.0 software (Plexon). Animals received pulse and sine wave stimulations according to their recording schedule (see below). Virally transduced neurons in reuniens were optically stimulated in blocks of 5 minutes (min, ON-blocks) with blue light (465 nm) using 60 ms ON-pulses at 1, 2, 4, 8 Hz frequencies, and 20 ms ON-pulses at 50 Hz across three recording sessions. On separate days, continuous sine wave stimulations were delivered in 5 min blocks at 1, 2, 4, 8 and 20 Hz frequencies. In all recording sessions, the first 5 min of recording (baseline) and the 5 min periods after each stimulation block were not optically stimulated. These OFF-blocks prevented the overheating of the brain tissue, and allowed activated neurons to return to baseline cell activity before the start of the next stimulation block.

#### Reuniens Optogenetic Recordings

Seven days prior to the first recording session, we habituated rats to the OF arena and recording equipment. During habituation days, rats were allowed to freely explore the OF for 20 min and restrained lightly to connect them from the tethered and optic patch cable. On the last 2 days of habituation, we familiarized rats to food treats (each treat: ∼1/8 fruit Loops) which were thrown from above the maze every 5 minutes to encourage rats to move around the arena. Neural data was acquired with a Plexon OmniplexD recording system (wide band: 40 kHz) and the PlexControl data acquisition software via six implanted 50 micron Teflon-coated SS electrode wires connected to an electrode interface board (6 channels; Neuralynx). LFP were sampled at 1 kHz.

#### Histology

After completion of all experiments, rats were anesthetized with isoflurane (5%) mixed with oxygen (800 ml/min) and marking lesions were made with a NanoZ (Plexon) to deliver 12 amperes for 10 s at each of the electrode locations. Rats were then transcardially perfused with 100 ml phosphate-buffered saline (PBS), followed by 200 ml of 4% paraformaldehyde (PFA, pH 7.4; Sigma-Aldrich). Brains were post-fixed overnight in 4% PFA and then placed in a 30% sucrose solution for cryoprotection. Frozen brains were cut on a Leica CM3050 S cryostat (40 mm; coronal plane) into three sets of immediately adjacent sections. One set was mounted, Nissl-stained, and coverslipped with Permount to visualize marking lesions. Marking lesions were then mapped onto plates of the Paxinos and Watson (2006).

In optogenetic subjects, an additional set of sections was incubated at room temperature for 48 hours with anti-green fluorescent protein (GFP) rabbit primary antibody (1:500, Rockland). After washes, tissue was incubated for 6 hours at room temperature in VectaFlour DyLight 488 anti-rabbit secondary. After incubation, tissue was washed in 0.1 M PB (3 × 5 min) then mounted on gelatin coated slides and coverslipped with VectaShield mounting medium with DAPI for visualization.

#### Quantification and Statistical Analysis Sequence Memory Performance Analysis

Performance on the task can be analyzed using a number of measures (Allen *et al*., 2014; Jayachandran *et al*., 2022). The first position of each sequence was excluded from all analysis as these items are always InSeq. Expected vs. observed frequencies were analyzed with G tests to determine whether the observed frequencies of InSeq and OutSeq responses for a given session were significantly different from the frequency expected by chance (see Table 1). G tests provide a measure of performance that controls for response bias and is a robust alternative to the χ^2^ test, especially for datasets that include cells with smaller frequencies (Sokal, 1995). To compare performance across sessions or animals, we calculated a sequence memory index as shown in the following equation:

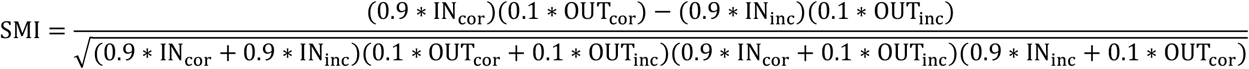

In essence, the SMI normalizes the proportion of InSeq and OutSeq items presented during a session and reduces sequence memory performance to a single value ranging from −1 to 1. A score of 1 represents perfect sequence memory, in which a subject would have correctly held his nosepoke response on all InSeq items and correctly withdrawn on all OutSeq items. A score of 0 indicates chance performance, such as if subjects responded to InSeq and OutSeq items with the same response pattern (e.g., holding until the signal 80% of the time regardless of the trial type). Negative SMI scores represent performance levels below that expected by chance.

#### LFP Analyses

We used custom MATLAB scripts with native signal processing functions (e.g ‘mscohere’, ‘trapz’,’envelope’), the Chronux toolbox (http://chronux.org/; Bokil *et al*., 2010), and circ_stats toolboxes for all analysis. We plotted spectrograms for each sessions and aligned to various timestamps (poke out for sequence task and stimulation events for optogenetic blocks) to observe the different frequencies during relevant events. Perievent spectrograms and coherograms were used to visualize and quantify LFP activity using custom MATLAB scripts. We focused our analyses on the 1-4 Hz (delta), 6–12 Hz (theta), and 15–30 Hz (beta), as our spectrograms showed high power in those bands at the level individual rats, and group. We compared across animals by *z*-normalizing values to calculate the t-score in spectrograms, and area under the curve (AUC) for coherence. We used traditional (parametric) *t* tests and ANOVAs for behavioral and frequency-based statistical comparisons. Tests were considered significant at *p* < 0.05.

#### Behavioral Tracking

Behaviorally relevant events were identified through behavioral tracking using DeepLabCut (DLC; Mathis *et al*., 2018) for sequence memory experiments. Markers were placed manually on the rodent’s nose, left and right ears, center of body/mass, tail base, and mid-tail. Markers were tagged on 200-350 (per experiment) on frames extracted from the recording video sessions to create a training dataset, and used to train the network using iterations. After initial training, output video files containing the continued markers and skeleton connecting structure were reviewed to identify out of place markers. Sample markers that had a likelihood below 0.90 were refined until they met criteria.

The exploratory behaviors (running and stationary periods) of rats in the sequence memory linear track were identified from the ‘x’ and ‘y’ markers’ coordinates per frame (DLC output) using MATLAB. These bouts were obtained using the exclusionary criteria defined by Schultheiss *et al*., (2020), based on previous studies observing locomotion in rodents (Maurer *et al*., 2005; McNaughton *et al*., 2006; Dombeck *et al*., 2007). For running bouts, bout minimum duration was specified as 2.05 s, and speeds to be above 2.5 cm/s (achieved using a median filter). For stationary bouts, the duration was also at least 2.05 s, with speeds below 2.5 cm/s (limited for linear track). Behaviors such as grooming and rearing were excluded using exclusionary criteria.

#### Beta Rhythm Bursts Detection

An envelope analysis was done on the raw LFP data to obtain *z*-scored upper envelopes (curves for the boundaries of the waveform) for all trials, which were averaged for each session. The raw data and *z*-scored envelopes were visually inspected for beta rhythm bursts across trials. A set of exclusionary parameters were defined based on this inspection to analyze beta bursts that met clear bursting criteria. Bursts were defined by minimum onset and offset amplitude values of 0.15 in the *z*-scored envelope, with a minimum peak in amplitude of 1.30 between a burst’s start and end. These were further separated from bursts that lasted under 0.95ms in order to consider bursts with at least two complete cycles in the raw waveform. Averages of the time to the first bursts’ onsets (latency) across trials, sessions, and subjects were calculated, as well as average durations for bursts in each region. Similarly, a cumulative density function (CDF) was used to compare the beta onsets between regions (co-occurrences of bursts).

