## Supplemental Figures for "Reuniens transiently synchronizes memory networks at beta frequencies"

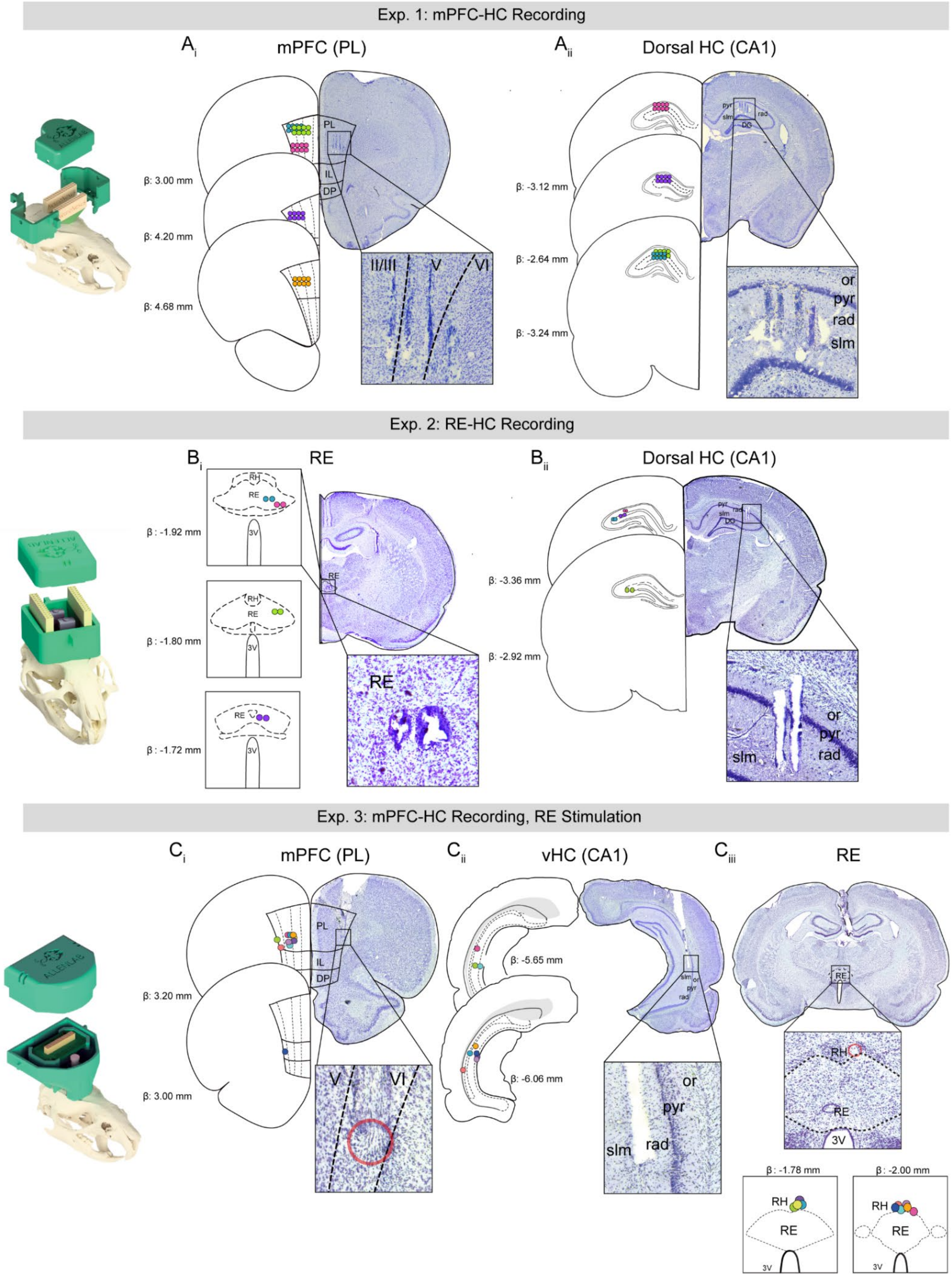

**Supplementary Figure 1: Electrode placements for all rats, Related to Figures 1, 2, and 3. A)** Representative nissl-stained images are shown for rats used in Experiment 1 with probes targeting **$A_{i}$)** mPFC and **$A_{ii}$)** dHC. Probes were implanted on the right side. Marking lesions were located across layers in prelimibic cortex and slm/rad. On the left we show a schematic representation of the targeted region and probe placement for all rats. Each rat is represented by a different colored circle indicating the exact locations. **B)** Representative nissl-stained image is shown for rats used in Experiment 2 with probes targeting **$B_{i}$)** RE and **$B_{ii}$)** dHC. Probes were implanted on the right side and the zoomed-in image shows that the marking lesions were located towards the lateral aspects of RE and slm/rad in CA1. On the left we show a schematic representation of the targeted region and probe placement for all rats. Each rat is represented by a different colored circle where we indicate the exact location. **C)** Representative nissl-stained image is shown for rats used in Experiment 3 with ss targeting **$C_{i}$)** mPFC and **$C_{ii}$)** vHC and **$C_{iii}$)** optrode placement in RE. We show a schematic representation of the targeted region and probe placement for all rats. Each rat is represented by a different colored circle indicating the exact locations. Abbreviations: mPFC, medial prefrontal cortex; HC, hippocampus; dHC, dorsal hippocampus; vHC, ventral hippocampus; RE, nucleus reuniens; RH, rhomboid nucleus; slm, stratum lacunosum-moleculare; rad, radiatum; pyr, pyramidal; or, stratum oriens; CA1, Cornus Ammonis 1; PL, prelimbic; IL, infralimbic; DP, dorasl peduncular cortex; ss, saggital sinus.

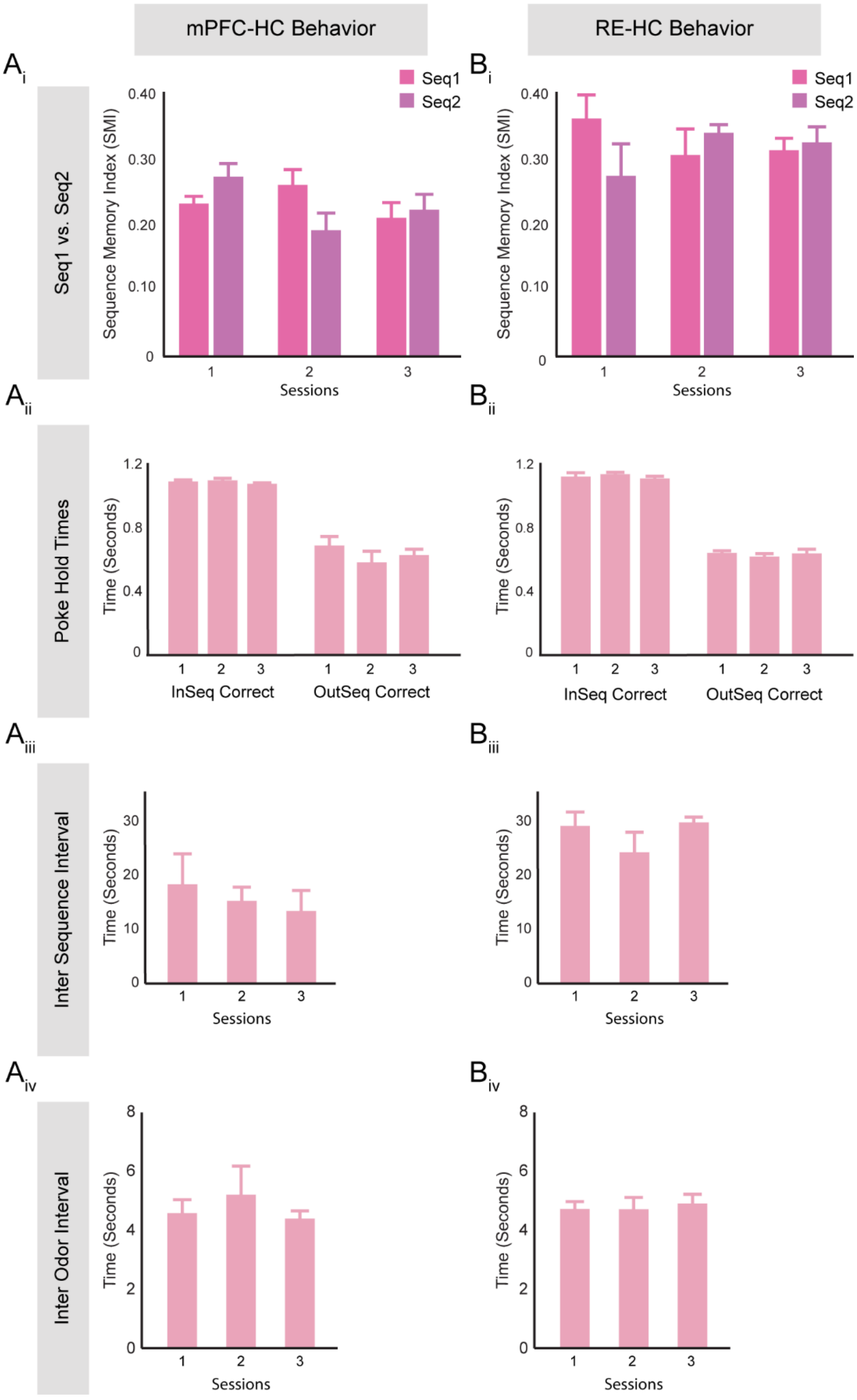
mPFC-HC Behavior
RE-HC Behavior
Ai
Bi
Seq1 vs. Seq2
Seq1
Seq2
Sequence Memory Index (SMI)
0.40
0.30
0.20
0.10
0
1
2
3
Sessions
Aii
Bii
Poke Hold Times
Time (Seconds)
1.2
0.8
0.4
InSeq Correct
OutSeq Correct
Aiii
Biii
Inter Sequence Interval
30
20
10
Aiv
Biv
Inter Odor Interval
8
6
4
2

**Supplementary Figure 2: Sequence memory and non-mnemonic aspects of behavior did not differ across sessions, Related to Figure 1 and 2. A)** Rat behavior on the sequence memory task used in the mPFC-HC experiment. **$A_i$)** SMI were not significantly different across sessions ($F_{(2,12)} = 2.40$, $p = 0.13$). Moreover, no significant differences between sequence 1 and sequence 2 across three sessions were found (Session 1: $t_{(4)} = -1.62$, $p = 0.14$; Session 2: $t_{(4)} = 1.85$, $p = 0.10$; Session 3: $t_{(4)} = -0.30$, $p = 0.77$). **$A_{ii}$)** There was no significant difference across the three sessions for $InSeq_{correct}$ poke times ($F_{(2,12)} = 0.34$, $p = 0.72$) and $OutSeq_{correct}$ poke times($F_{(2,12)} = 0.42$, $p = 0.67$), indicating that nose-poke behavior was not affected by sessions. **$A_{iii}$)** ISI did not differ significantly across sessions ($F_{(2,12)} = 0.96$, $p = 0.41$), suggesting rats in all three conditions ran at similar rates between sequences. **$A_{iv}$)** IOI did not significantly differ across sessions ($F_{(2,12)} = 0.62$, $p = 0.55$), suggesting rats collected water rewards and engaged odors at similar rates. **B)** Rats' behavior on the sequence memory task used in the RE-HC experiment. **$B_i$)** SMI were not significantly different across sessions ($F_{(2,11)} = 0.04$, $p = 0.96$). We found no significant differences between sequence 1 and sequence 2 over the three sessions (Session 1: $t_{(3)} = 1.75$, $p = 0.18$; Session 2: $t_{(3)} = -1.09$, $p = 0.36$; Session 3: $t_{(3)} = -0.51$, $p = 0.65$). **$B_{ii}$)** There was no significant difference across sessions for $InSeq_{correct}$ poke times ($F_{(2,11)} = 0.79$, $p = 0.49$) and $OutSeq_{correct}$ poke times ($F_{(2,11)} = 0.44$, $p = 0.66$), indicating that nose-poke behavior was not affected by session. **$B_{iii}$)** ISI did not differ significantly between sessions ($F_{(2,11)} = 1.41$, $p = 0.29$). **$B_{iv}$)** IOI did not differ significantly across sessions ($F_{(2,11)} = 0.12$, $p = 0.89$). Abbreviations: mPFC, medial prefrontal cortex; HC, hippocampus; $InSeq_{correct}$, in-sequence correct; $OutSeq_{correct}$, out-of-sequence correct; ISI, inter sequence interval; IOI, inter odor interval; SMI, sequence memory index.

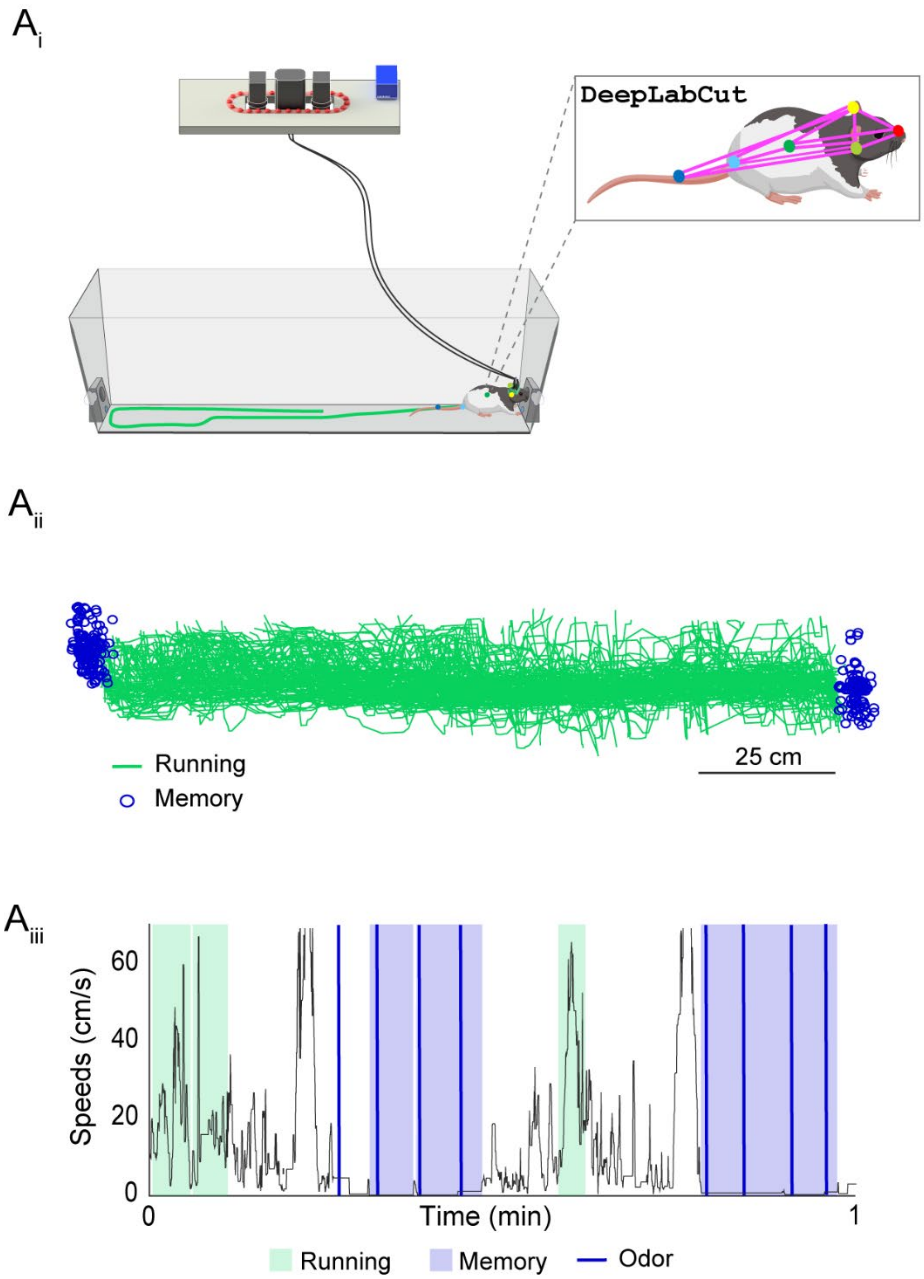


**Supplementary Figure 3: Behavioral conditions obtained using DeepLabCut for sequence memory task for separation in electrophysiological analyses, Related to Figure 1, 2, and 4.** Behavioral tracking was done using a deep machine learning algorithm from training datasets to achieve quantifiable and accurate animal pose estimation ([www.mackenziemathislab.org/deeplabcut](www.mackenziemathislab.org/deeplabcut)). This approximation was further filtered using custom written scripts in MATLAB. **$A_i$)** Rat in sequence memory task with behavioral tracking markers for trained and learned body parts (nose, left and right ears, center of gravity, tail start, and middle of tail; inset). **$A_{ii}$)** Representative processed trajectories from one rat showing active running in between odors (represented in green), and odor sampling (memory) occurring during stationary positions (represented in blue). **$A_{iii}$)** Sample activity plot (~1 min) with filtered speeds during the sequence memory task. Running and stationary bouts are highlighted in green and blue respectively, as well as odor trial times (blue lines).

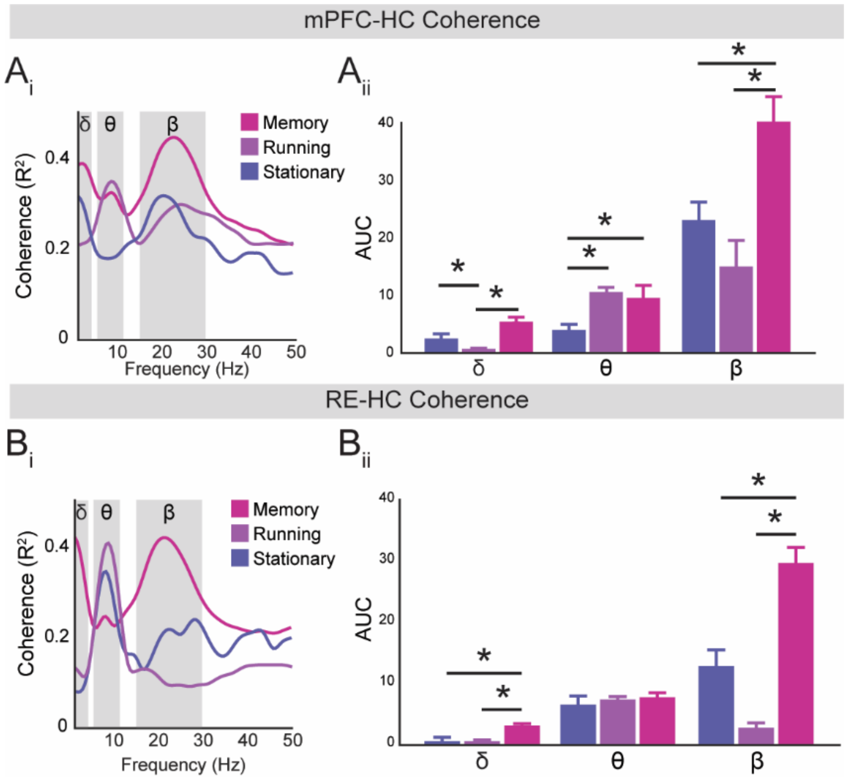


**Supplementary Figure 4: Delta, theta, and beta during different behavioral conditions, Related to Figure 1 and 2. $A_i$)** mPFC-HC coherence showing memory, running, and stationary periods. During memory both delta and beta increase while theta remains the same. During stationary periods (separate from stationary during the sequence trials) theta decreases while delta and beta increase in coherence, and all three frequency bands had lower coherence compared to memory periods. Finally, during running theta coherence increased while delta and beta decreased. **$A_{ii}$)** Delta was significantly different between the three behavioral conditions ($F_{(2,11)}$ = 12.46, $p$ = 3.0x10$^{-3}$). Theta was significantly different between the three behavioral conditions ($F_{(2,11)}$ = 6.96, $p$ = 0.02). Beta was significantly different between the three behavioral conditions ($F_{(2,11)}$ 8.38, $p$ = 0.01). **$B_i$)** RE-HC coherence showing memory, running, and stationary periods. During memory, both delta and beta increased while theta decreased. During stationary periods (separate from stationary during the sequence trials) theta increased while beta had a moderate increase in coherence. Finally, during running theta coherence increased while delta and beta decreased. **$B_{ii}$)** Delta was significantly different between the three behavioral conditions ($F_{(2,11)}$ =11.52, $p$ = 3.0x10$^{-3}$). Theta was not significant different between the three behavioral conditions ($F_{(2,11)}$ = 0.87, $p$ = 0.45). Beta was significantly different between the three behavioral conditions ($F_{(2,11)}$ 43.68, $p$ = 2.3x10$^{-5}$). Abbreviations: mPFC, medial prefrontal cortex; HC, hippocampus; RE, nucleus reuniens; δ, delta; θ, theta; β, beta.

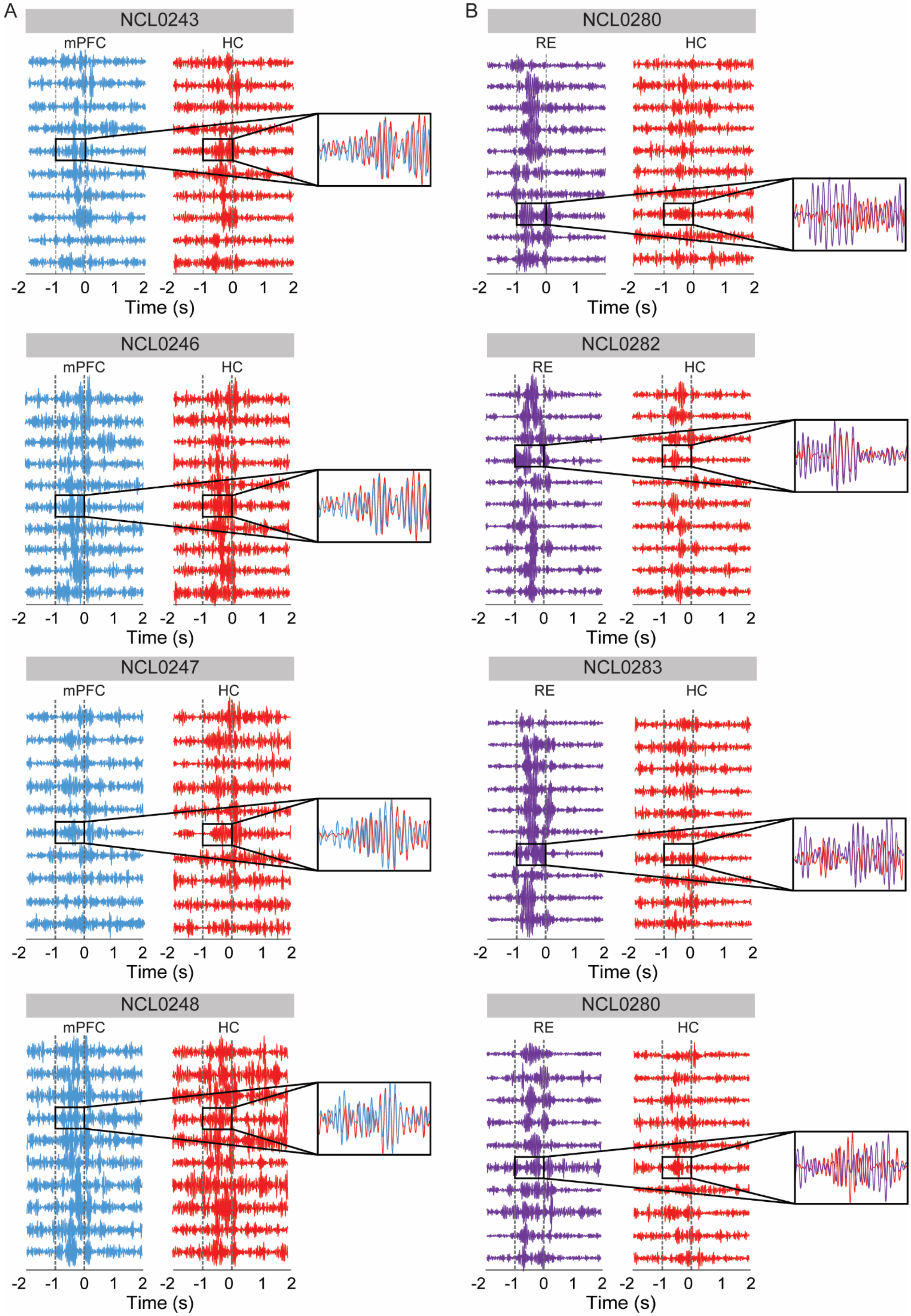
A
NCL0243
mPFC
HC
-2 -1 0 1 2 -2 -1 0 1 2
Time (s)
NCL0246
mPFC
HC
-2 -1 0 1 2 -2 -1 0 1 2
Time (s)
NCL0247
mPFC
HC
-2 -1 0 1 2 -2 -1 0 1 2
Time (s)
NCL0248
mPFC
HC
-2 -1 0 1 2 -2 -1 0 1 2
Time (s)
B
NCL0280
RE
HC
-2 -1 0 1 -2 -1 0 1 2
Time (s)
NCL0282
RE
HC
-2 -1 0 1 2 -2 -1 0 1 2
Time (s)
NCL0283
RE
HC
-2 -1 0 1 2 -2 -1 0 1 2
Time (s)
NCL0280
RE
HC
-2 -1 0 1 2 -2 -1 0 1 2
Time (s)

**Supplementary Figure 5. Sample beta-filtered voltage traces during sequence trials, Related to Figure 1 and 2.** Bandpass filtered beta during sequence trials for a subsample of InSeq trials across rats in both groups (mPFC-HC and RE-HC). **A)** Prefrontal cortex and hippocampus showed clear beta synchorny across trials in all rats. **B)** Reuniens beta bursts occured earlier and had a larger amplitdue compared to hippocampus. Abbreviations: InSeq, in-sequence; mPFC, medial prefrontal fortex; HC, hippocampus; RE, nucleus reuniens; NCLXXXX, rat ID.

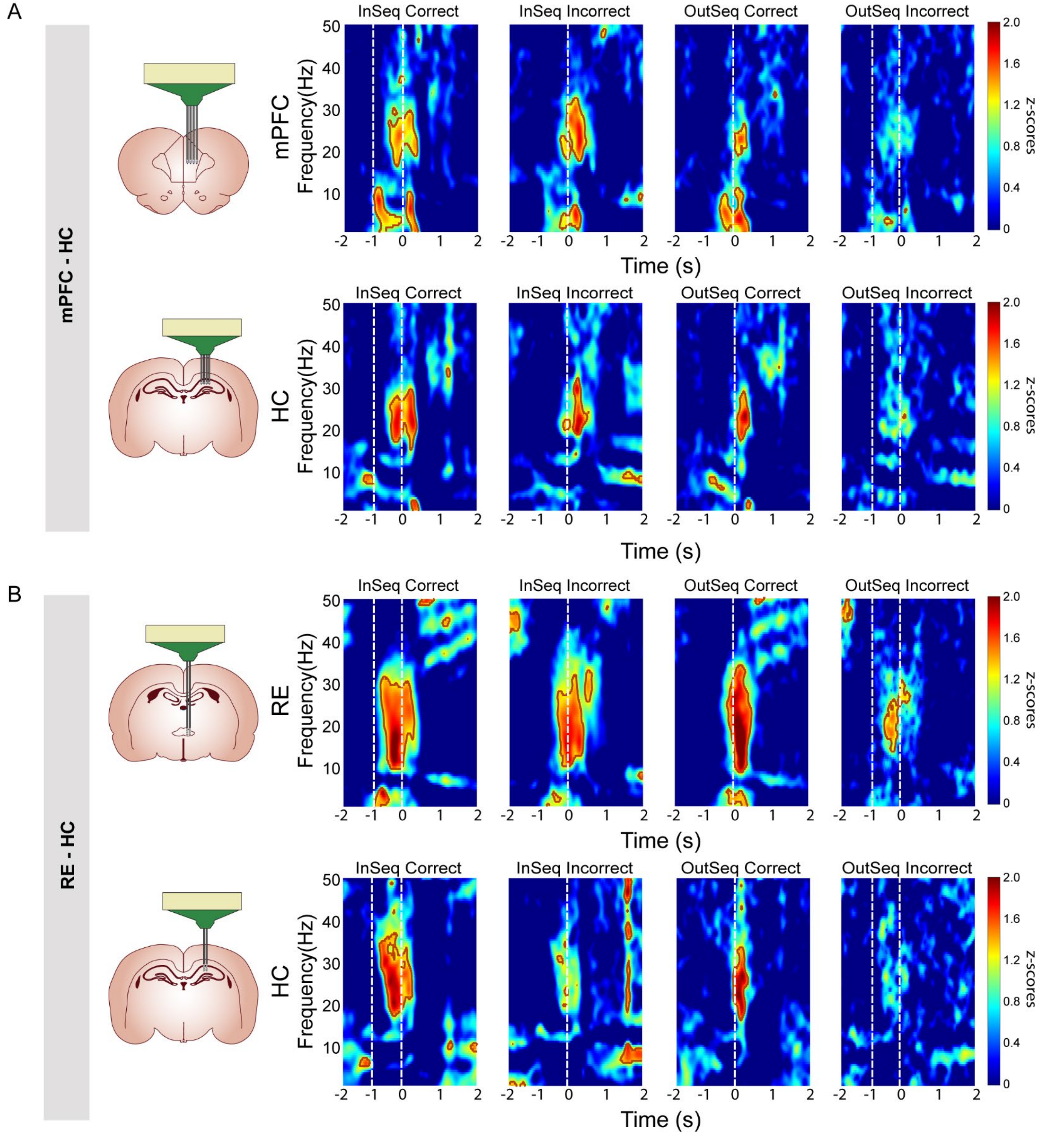


**Supplementary Figure 6: Prefrontal cortex, reuniens, and hippocampus perievent spectrograms during the sequence memory task, Related to Figure 1 and 2. A)** mPFC and HC PESGs across different trial types aligned to poke out. Beta can be seen in both mPFC and HC. The significant areas (z-tests) are outlined where $p < 0.05$ (brick color). OutSeq trials did not show significant beta in mPFC or HC rats. **B)** RE and HC PESGs across different trial types aligned to poke out. Beta can be seen in both RE and HC. The significant areas are outlined where $p < 0.05$ (brick color). HC OutSeq incorrect trials failed to show significant beta. Abbreviations: PESG, perievent spectrogram; mPFC, medial prefrontal cortex; HC, hippocampus; RE, nucleus reuniens; InSeq, in sequence; OutSeq, out of sequence; $\text{InSeq}_{\text{correct}}$, in-sequence correct, $\text{InSeq}_{\text{incorrect}}$, in-sequence incorrect, $\text{OutSeq}_{\text{correct}}$, out-of-sequence correct, $\text{OutSeq}_{\text{incorrect}}$, out-of-sequence incorrect.

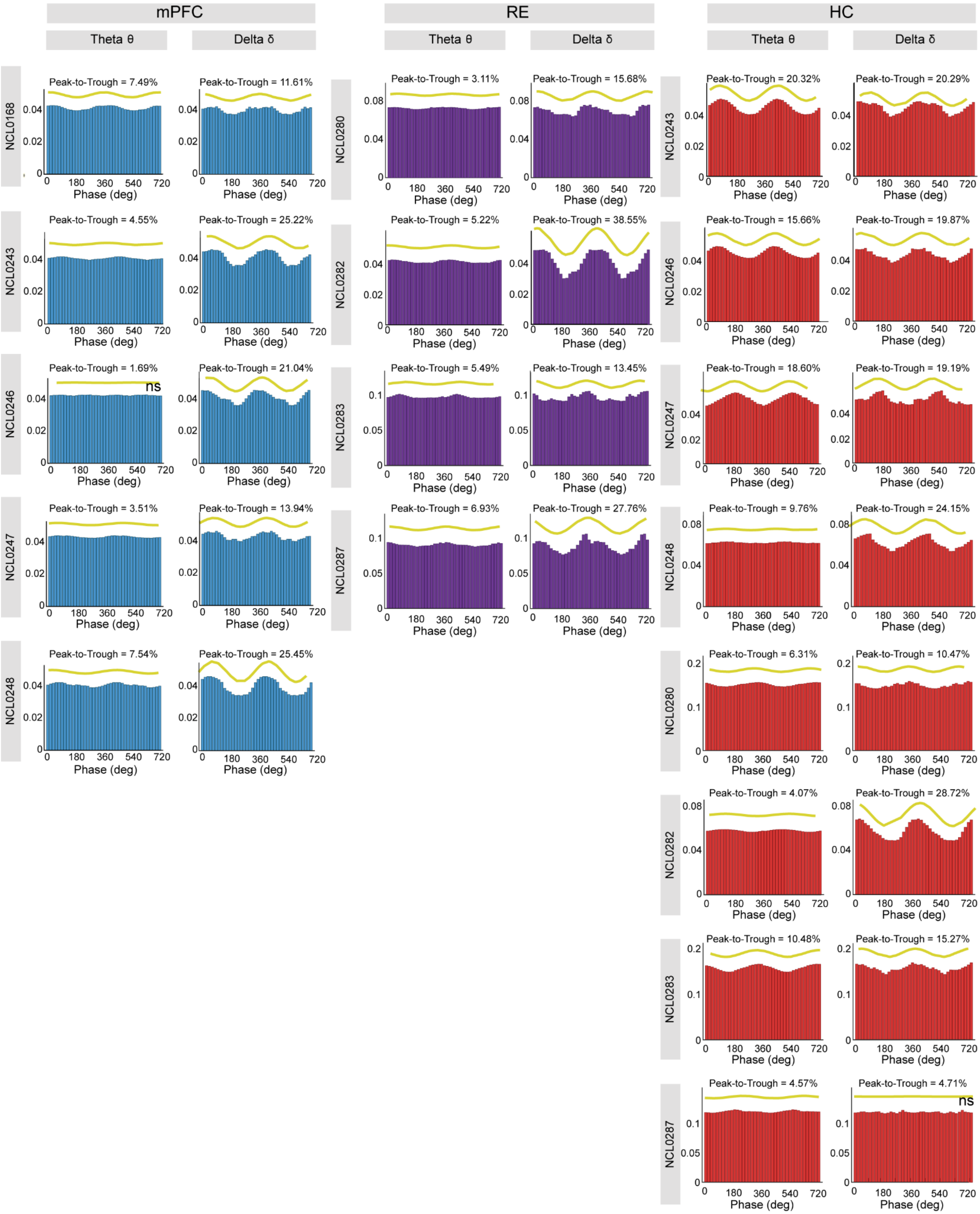
mPFC
RE
HC
Theta θ
Delta δ
NCL0168
Peak-to-Trough = 7.49%
Peak-to-Trough = 11.61%
NCL0243
Peak-to-Trough = 4.55%
Peak-to-Trough = 25.22%
NCL0246
Peak-to-Trough = 1.69%
ns
Peak-to-Trough = 21.04%
NCL0247
Peak-to-Trough = 3.51%
Peak-to-Trough = 13.94%
NCL0248
Peak-to-Trough = 7.54%
Peak-to-Trough = 25.45%
NCL0280
Peak-to-Trough = 3.11%
Peak-to-Trough = 15.68%
NCL0282
Peak-to-Trough = 5.22%
Peak-to-Trough = 38.55%
NCL0283
Peak-to-Trough = 5.49%
Peak-to-Trough = 13.45%
NCL0287
Peak-to-Trough = 6.93%
Peak-to-Trough = 27.76%
NCL0243
Peak-to-Trough = 20.32%
Peak-to-Trough = 20.29%
NCL0246
Peak-to-Trough = 15.66%
Peak-to-Trough = 19.87%
NCL0247
Peak-to-Trough = 18.60%
Peak-to-Trough = 19.19%
NCL0248
Peak-to-Trough = 9.76%
Peak-to-Trough = 24.15%
NCL0280
Peak-to-Trough = 6.31%
Peak-to-Trough = 10.47%
NCL0282
Peak-to-Trough = 4.07%
Peak-to-Trough = 28.72%
NCL0283
Peak-to-Trough = 10.48%
Peak-to-Trough = 15.27%
NCL0287
Peak-to-Trough = 4.57%
Peak-to-Trough = 4.71%
ns
Phase (deg)

**Supplementary Figure 7. Average beta amplitude-phase coupling with theta and delta during sequence memory trials, Related to Figure 3.** Phase-amplitude plots showing the mean beta amplitudes along the phase of theta or delta rhythms recorded on the same electrodes. Subjects were averaged across sessions within regions of interest (prefrontal cortex, blue; reuniens, purple; hippocampus, red). Bar graphs show magnitude of beta amplitude at a phase degree with a fitted sine wave (yellow) and calculated percentage peak-to-trough distance on top. All subjects were found to have a significant relationship between fitted sine wave and plotted data with $p < 0.01$, except for two subjects noted above with ns. Abbreviations: LFP, local field potential; ns, not significant; NCLXXXX, rat ID.

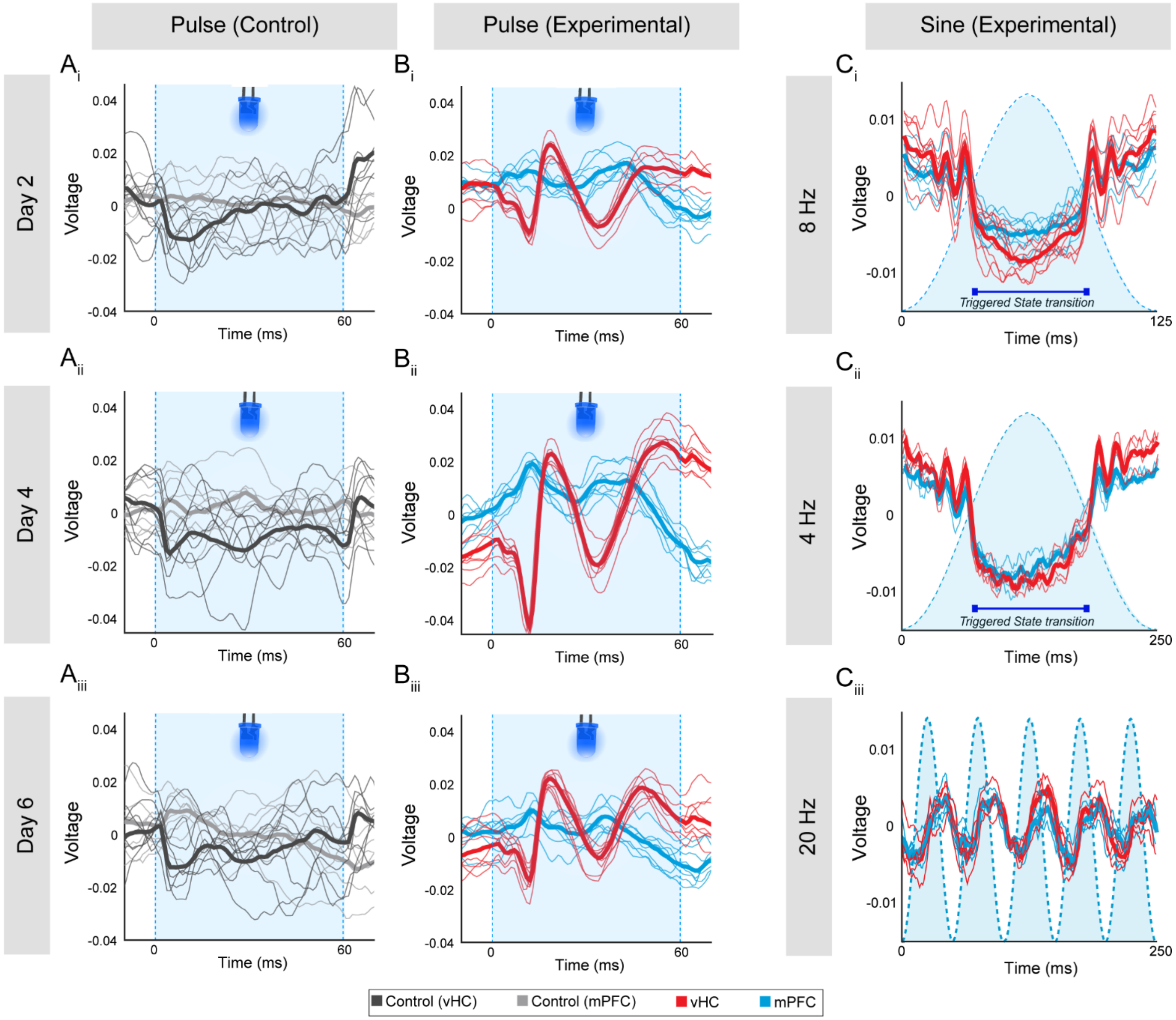


**Supplementary Figure 8: Sample reuniens blue light evoked responses from ventral hippocampus, Related to Figure 4.** Stacked mPFC (blue) and vCA1 (red) raw signals from an averaged one second period 8 Hz blue light pulse stimulation (60 ms each) during the first session (day 2), second (day 4) and third session (day 6) in viral controls $\mathbf{A_i}$**-**$\mathbf{A_{iii}}$**)**, and experimental (ChR2) subjects $\mathbf{B_i}$**-**$\mathbf{B_{iii}}$**)**. Thick lines represent the mean raw signal across all pulses and thinner lines represent raw signal during each pulse. Note that in the controls the activity remains relative the same across pulses and days. In contrast, in the ChR2, a very strong evoked vCA1 monosynaptic responses (red) are observed consistently between pulses and across days. A similar but less strong response can be observed in mPFC (light blue). **C)** Sine wave stimulations also changed LFP voltage activity in mPFC (blue) and vCA1 (red) in ChR2 rats. Low ($\mathbf{C_{i\text{-}ii}}$) and high ($\mathbf{C_{iii}}$) sine waves triggered rapid down-state transitions at the rise of the sine wave stimulus, and up-state transitions with the fall of the sine that created rhythmic and phased locked voltage activity in both brain regions. This demonstrated that optogenetic excitation of RE neurons effectively elicited electrophysiological responses in mPFC and hippocampus. Shaded light blue area demonstrates the duration of the blue light pulse. Abbreviations: mPFC, medial prefrontal cortex; vHC, ventral hippocampus; RE, nucleus reuniens; :FP, local field potential; vCA1, ventral Cornus Ammonis 1; ChR2, channelrhodopsin.

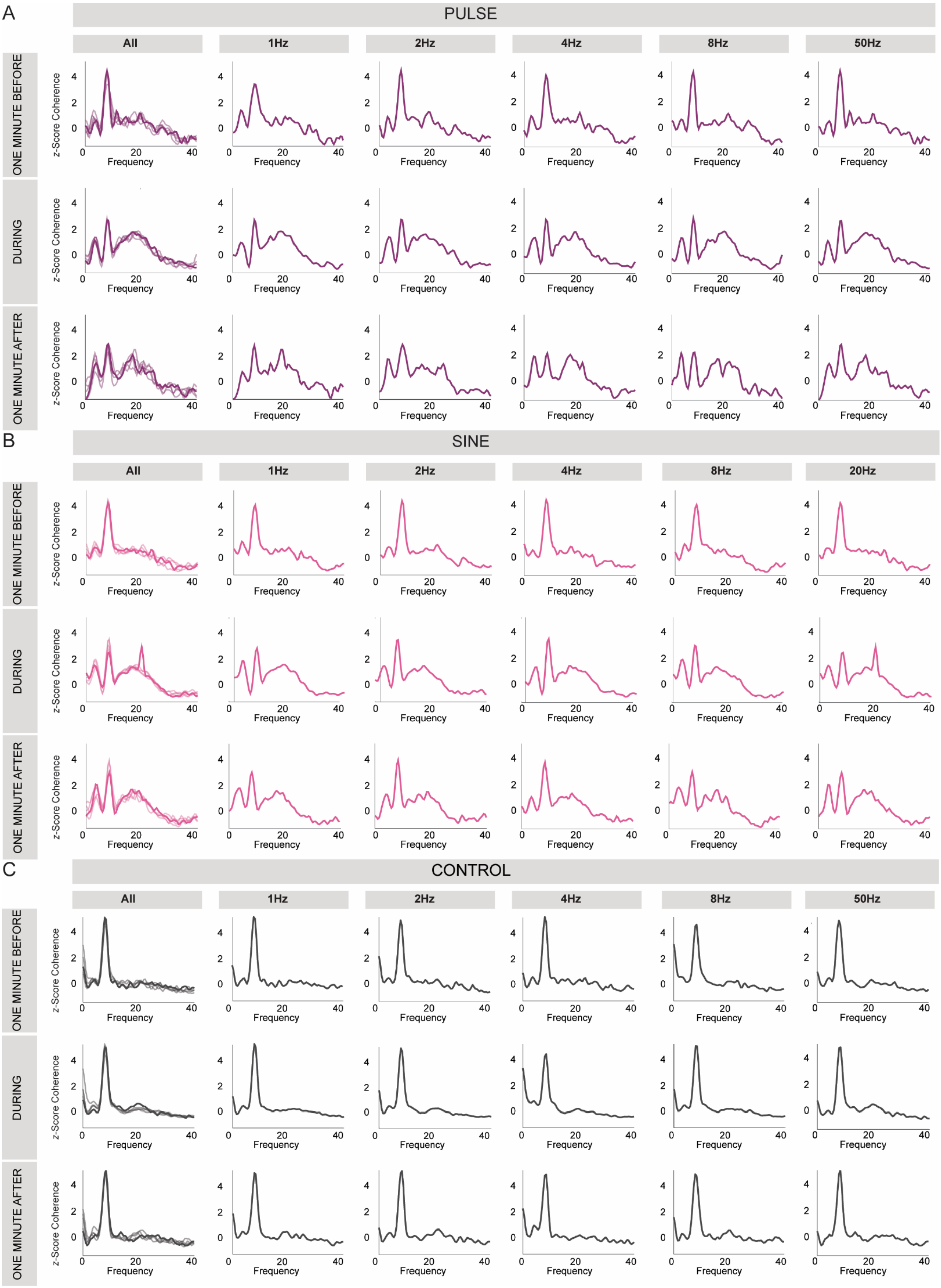
A
PULSE
All
1Hz
2Hz
4Hz
8Hz
50Hz
ONE MINUTE BEFORE
DURING
ONE MINUTE AFTER
z-Score Coherence
Frequency
B
SINE
All
1Hz
2Hz
4Hz
8Hz
20Hz
ONE MINUTE BEFORE
DURING
ONE MINUTE AFTER
C
CONTROL
All
1Hz
2Hz
4Hz
8Hz
50Hz
ONE MINUTE BEFORE
DURING
ONE MINUTE AFTER

**Supplementary Figure 9. Prefrontal-hippocampal coherence curves were similar during and after reuniens activations regardless of the frequency stimulation applied, Related to Figure 4.** mPFC-HC coherence was assessed one minute before, during and one minute after RE optogenetic stimulations. Pulse **(A)** and sine **(B)** stimulation patterns showed nearly identical coherence curves (when stacked on top of each other—see All columns) during and after stimulations regardless of the frequency (1, 2, 4, 8, 50 or 20 Hz) with prominent increases in the delta and beta bands and decreases in the theta band. **C)** In control rats (across all time points), the coherence curves were theta dominated and totally unaffected by pulse and sine stimulation patterns, as expected. Coherence curves are shown one minute before stimulations with all showing a predominant theta coherence. Abbreviations: mPFC, medial prefrontal cortex; HC, hippocampus; RE, nucleus reuniens; ChR2, channelrhodopsin.
